# Desmosomes polarize mechanical signaling to govern epidermal tissue form and function

**DOI:** 10.1101/2020.01.21.914176

**Authors:** Joshua A. Broussard, Jennifer L. Koetsier, Marihan Hegazy, Kathleen J. Green

## Abstract

The epidermis is a stratified epithelium in which structural and functional features are polarized across multiple cell layers. This type of polarity is essential for establishing the epidermal barrier, but how it is created and sustained is poorly understood. Previous work identified a role for the classical cadherin/filamentous-actin network in establishment of epidermal polarity. However, little is known about potential roles of the most prominent epidermal intercellular junction, the desmosome, in establishing epidermal polarity, in spite of the fact that desmosome constituents are patterned across the apical to basal cell layers. Here, we show that desmosomes and their associated intermediate filaments (IF) are key regulators of mechanical polarization in epidermis, whereby basal and suprabasal cells experience different forces that drive layer-specific functions. Uncoupling desmosomes and IF or specific targeting of apical desmosomes through depletion of the superficial desmosomal cadherin, desmoglein 1, impedes basal stratification in an in vitro competition assay and suprabasal tight junction barrier functions in 3D reconstructed epidermis. Surprisingly, disengaging desmosomes from IF also accelerated the expression of differentiation markers, through precocious activation of the mechanosensitive transcriptional regulator serum response factor (SRF) and downstream activation of Epidermal Growth Factor Receptor family member ErbB2 by Src family kinase (SFK) mediated phosphorylation. This Dsg1-SFK-ErbB2 axis also helps maintain tight junctions and barrier function later in differentiation. Together, these data demonstrate that the desmosome-IF network is a critical contributor to the cytoskeletal-adhesive machinery that supports the polarized function of the epidermis.

## Introduction

To transition from single celled organisms to multicellular metazoans, cells evolved multiple mechanisms to form stabilized cell-cell interactions [1]. Ultimately, this process led to the development of simple adherent epithelial sheets, which serve as a protective barrier between an organism and its environment. Additional epithelial and junctional complexity arose later in evolution, with the emergence of multilayered, stratified epithelia, including elaborate tissues like the epidermis [2]. In vertebrates, this multilayered epithelium serves as a barrier against water loss, mechanical insults, and as a physical and immune barrier to external pathogens (containing both innate and adaptive immune elements) [3, 4]. The epidermis comprises four distinct layers, basal, spinous, granular (SG), and cornified, which undergo constant regeneration. Regeneration occurs through a process whereby keratinocytes in the basal layer proliferate, at some point commit to a program of terminal differentiation, and exit the basal layer. During differentiation, cells maintain cell-cell interactions while moving progressively into the suprabasal layers where they are incorporated into the epidermal barrier and ultimately slough off of the skin’s surface.

Polarity, the differential patterning of structural and functional features along an axis, is a fundamental property of all epithelial tissues and is essential to their function. Whereas simple epithelia exhibit apical to basal (apicobasal) polarization within a single cell layer, the epidermis exhibits polarization across multiple cell layers, through poorly understood mechanisms [2]. For example, barrier-essential tight junctions (TJs) are found in association with adherens junctions on the apicolateral surface of simple epithelia. In the epidermis, functional TJs localize in association with adherens junctions to the second of three SG layers [5]. Improper development or disturbed maintenance of the epidermal TJ barrier is associated with a variety of inflammatory skin diseases (e.g., atopic dermatitis and psoriasis), impaired wound healing, and cancer [6]. In addition, simple epithelia exhibit a polarization of intercellular forces, with a region of high tension generated by actomyosin near the apical surface [7]. This region of high tension can regulate tissue morphology, homeostasis, and barrier function [7]. Moreover, loss of polarized mechanical tension is associated with cancer morphogenesis in multiple types of epithelial tissues [8]. Recent work has also identified a stiffness gradient in both human and mouse epidermis, with progressive stiffening from basal to superficial layers [9]. However, how such a mechanical asymmetry contributes to polarized cell signaling and behavior is not well understood.

Cell-cell adhesion complexes and their associated cytoskeletal networks are prime candidates to regulate polarized tissue mechanics. Desmosomes are calcium-dependent, intercellular adhesions that link to the intermediate filament (IF) cytoskeleton. Desmosomes comprise three main protein families: transmembrane cadherins (desmogleins and desmocollins), armadillo proteins (plakophilins and plakoglobin), and plakin proteins (desmoplakin, DP). Cadherins of adjacent cells form trans-interactions to mediate cell-cell adhesion. Intracellularly, cadherins interact with armadillo proteins, which are coupled to IF through DP. In the epidermis, desmosomal cadherins display graded patterns of expression. Some, such as desmoglein 3 (Dsg3), are concentrated in the proliferating basal layer, while others, such as desmoglein 1 (Dsg1), are concentrated in the superficial layers. This is in contrast to the adherens junction component E-cadherin, which is expressed throughout the epidermis [10].

A role for filamentous-actin (F-actin)-based adhesive networks in regulating mechanics (tension and stiffness) in conjunction with adherens junctions has been well-established [11, 12]. However, the contributions of the desmosome/IF network in regulating cell mechanics are not as well understood. Our previous work elucidated a role for desmosomes in mechanical regulation by modulating the interaction between desmosomes and IF using mutant forms of DP in a single layered epithelial system. Through a combination of micropillar arrays and atomic force microscopy (AFM), we demonstrated that strengthening DP-IF interactions increased cell-cell tensional forces and cell stiffness, while disrupting the interaction decreased these forces [12]. Moreover, these effects required cooperation with F-actin-based networks, indicating that desmosome/IF and F-actin-based networks function synergistically to regulate cell mechanics, highlighting their integrated nature. How these integrated networks work together to drive morphogenetic events in more complex tissues is not well understood.

The patterned expression of the desmosome/IF network suggests their potential role in governing the differential biophysical properties of distinct epidermal layers. Here, we provide evidence that the desmosome/IF linkage governs the ability of the epidermis to adopt a highly polarized mechanical phenotype. Disruption of this linkage alters indicators of mechanical asymmetries and the normal polarized functions of the epidermis including stratification, differentiation, and the establishment of an intact epidermal barrier. Finally, we identify a putative, mechanically-sensitive serum response factor (SRF)-Src family kinase (SFK)-ErbB2 signaling axis through which the desmosome/IF linkage promotes these polarized epidermal functions.

## Results

### Uncoupling the desmosome/IF connection induces rearrangements of cytoskeletal/adhesive complexes in epidermal keratinocytes

To establish models for addressing the role of the desmosome/IF network in epithelia that have the ability to stratify, we performed loss-of-function experiments utilizing a well-characterized mutant of the cytolinker protein DP in two established submerged models of neonatal human epidermal keratinocytes (NHEKs).

DP provides the connection between the desmosome core components and the IF cytoskeleton (Figure 1A). This mutant, referred to as DPNTP (Figure 1A), consists of the first 584 amino acids of DP. It retains the ability to interact with the desmosomal core but lacks the IF-binding domains, acting as a dominant negative mutant to uncouple desmosomes from the IF network [12–14]. We previously showed that expression of the uncoupling mutant in a simple epithelial A431 cell model (an epidermoid cell line) results in retraction of the IF cytoskeleton from sites of cell-cell adhesion [12].

**Figure 1.**
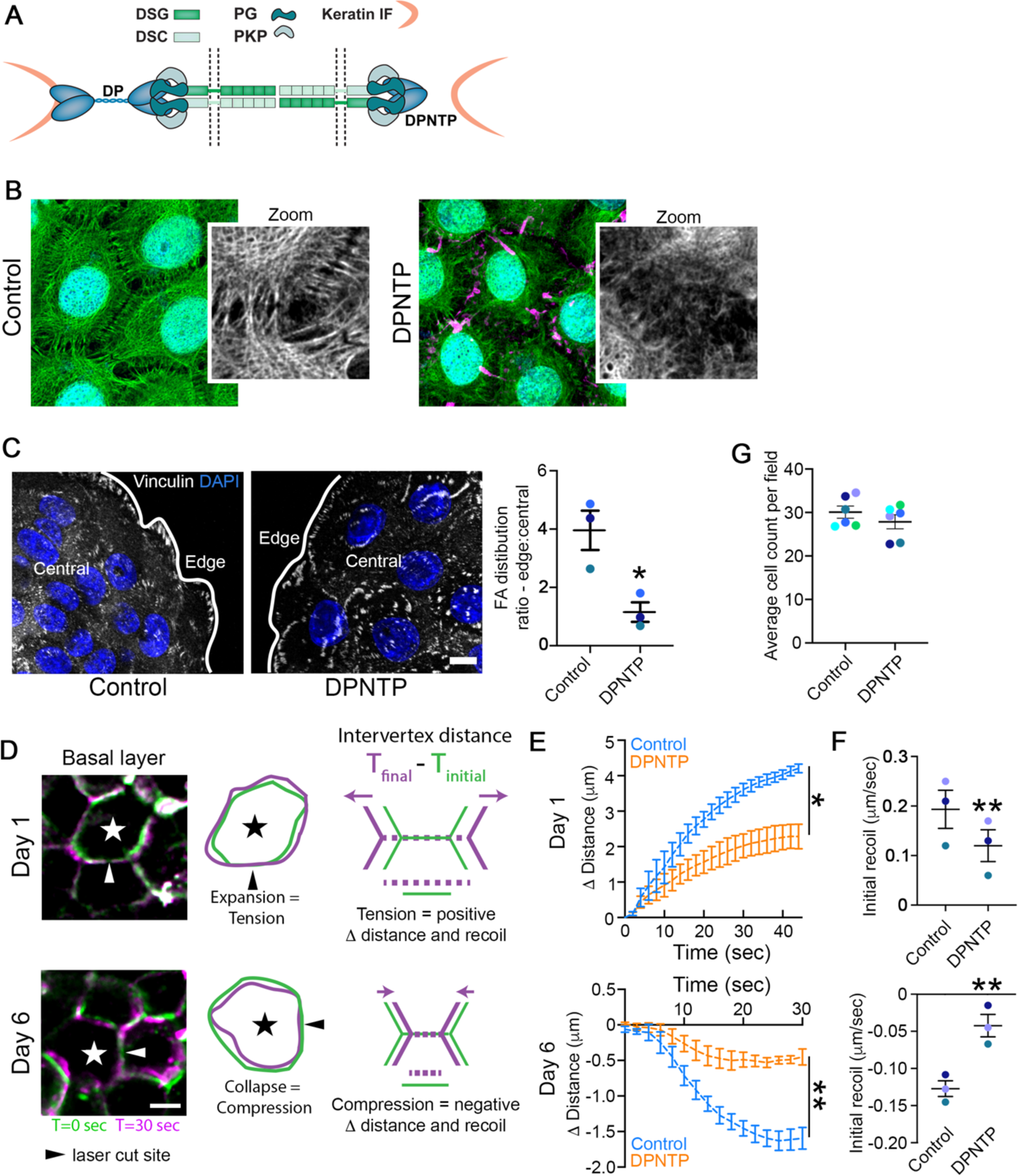
Uncoupling the desmosome/IF connection induces rearrangements of cytoskeletal/adhesive complexes. A) Schematic depicts the interactions among the desmosomal core proteins and the intermediate filament (IF) cytoskeleton in both a control condition (left) as well as when the desmosomal core is uncoupled from IF (right). DP, desmoplakin; PKP, plakophilin; PG, plakoglobin; DSG, desmoglein; DSC, desmocollin. B) Maximum projection micrographs show immunofluorescence staining of the keratin 5/14 (K5/14) IF cytoskeleton in basal cells of differentiating monolayers of NHEKs expressing either GFP (Control) or DPNTP-GFP. K5/14 is pseudocolored green and DPNTP is shown in magenta. DAPI indicates nuclei in blue and bar is 10 μm. C) Left, maximum projection micrographs show immunofluorescence staining of vinculin at the basal/substrate interface of differentiating NHEK ephrin colonies expressing either GFP (Control) or DPNTP-GFP. DAPI indicates nuclei in blue and bar is 10 μm. Right, the average vinculin-positive area per cell was quantified for both cells on the edge and in the central region of the colony. The average ratio (dashed line) of edge to central cells is shown for control and DPNTP-expressing colonies. *p=0.0233, paired t-test from three independent experiments, error bars are SEM. D) Images show the membranes (indicated by myr-tomato) of basal cells within epidermal equivalent cultures before and after laser ablation at the indicated times. At day 1, cells expand upon ablation, indicating tension within the layer and corresponding to a positive value for measured intervertex distance and initial recoil velocity. At day 6, cells collapse upon ablation, indicating compression within the layer and corresponding to a negative value for measured intervertex distance and initial recoil velocity. Bar is 5μm. E) Quantification is shown for the average change in intervertex distance (*Δ* Distance) over time for basal cells after ablation within control and DPNTP-expressing epidermal equivalent cultures both at day 1 and day 6 of their development. *p=0.0361, **p=0.0058, two-way ANOVA with repeated measures from three independent experiments, error bars are SEM. F) Quantification is shown for the average initial recoil velocity after ablation for basal cells within control and DPNTP-expressing epidermal equivalent cultures both at day 1 (upper graph) and day 6 (lower graph) of their development. **p<0.009, paired t-test from three independent experiments, error bar is SEM. G) The average cell count per field is shown for control and DPNTP-expressing epidermal equivalent cultures both at day 6 of their development. Dashed line is mean of 6 independent experiments, error bars are SEM.

We first expressed DPNTP tagged with GFP in monolayers of NHEKs induced to differentiate and stratify with high calcium containing medium (calcium switch) [15] (Figure S1A). After 2 days of calcium-induced cell-cell adhesion in NHEK monolayers, keratin IFs closely adjoin the cell-cell interface as they are anchored there by functional desmosomes (Figure 1B). Compared with cytoplasmic GFP controls, expression of DPNTP in NHEKs resulted in loss of orthogonally anchored keratin IF bundles, indicating the successful uncoupling of the desmosome/IF connection (Figure 1B).

In tissues, complex patterns of cell-cell and cell-substrate forces are balanced and both contribute to epithelial tissue morphogenesis [16]. We previously showed that uncoupling the desmosome/IF connection in A431 cells was accompanied by a significant reduction in cell-cell junctional forces [12]. Uncoupling the desmosome/IF linkage in ephrin-induced colonies of NHEK monolayers (Figure S1B) with DPNTP resulted in an altered distribution of cell-substrate contacts. In control colonies, vinculin-containing focal contacts are enriched near the colony edge (Figure 1C). However, uncoupling desmosomes/IF with DPNTP altered this staining pattern with vinculin-positive substrate contacts scattered throughout the basal surface of the colony, significantly decreasing the enrichment of focal adhesions at the colony edge (Figure 1C). These results suggested a mechanical cell-cell to cell-substrate switch [17, 18], which is consistent with previous reports in cell culture and in drosophila where loss of cell–cell adhesion increased cell–substrate adhesion [17–19].

### Uncoupling the desmosome/IF connection alters the mechanical properties of epidermal keratinocytes

Since mechanical forces regulate cell behavior, their polarized distribution may drive segregation of tissue functions. Tensile (pulling) and compressive (pushing) forces are known to affect the behavior of cells within epithelia, regulating cell division, migration, tissue morphogenesis, as well as promoting cancer cell invasion [20, 21]. In order to directly test the possibility that the desmosome/IF complex controls the mechanical properties of epidermal keratinocytes during development of the polarized tissue, we carried out laser ablation experiments in epidermal equivalent cultures (Figure S1C). Laser ablation was performed at early and late stages of in vitro morphogenesis to examine development of mechanical behaviors of basal cells over time (Figure 1D). Uncoupling the desmosome/IF linkage with DPNTP resulted in a significant reduction in recoil behavior at both days 1 and 6, indicating reduced cell-cell forces (Figure 1E and F). Interestingly, 6 days after lifting to the air-liquid interface, cells within the basal layer exhibited a collapsing behavior that would be consistent with cells experiencing compressive forces due to crowding and/or tissue jamming. Importantly, examination of the cell density indicated there was no significant effect of DPNTP expression on cell density (Figure 1G), suggesting the effects on mechanical behavior are not caused by altered forces due to crowding. These data are consistent with our previous observations that in A431 cells cell-cell junctional tugging forces and cortical stiffness was dependent on DP-IF interactions, working in concert with the actin cytoskeleton [12]. Thus, the idea that proper integration of DP-IF connections with other cytoskeletal networks is required for cooperative control of cell mechanics [22–24] also applies to developing complex epithelia.

### Uncoupling the desmosome/IF connection hinders epidermal keratinocyte stratification

In models of cell extrusion, monolayers of simple epithelial tissues maintain a constant cell density by eliminating cells that are normally fated to die. This process requires tight control of dynamic alterations in cell-cell mechanics and depends on actomyosin-generated forces [25–27]. In the case of the epidermis, cells undergo a somewhat similar process whereby they lose cell-substrate contacts as they move out of the basal layer [28], but once they reach the second layer they remain attached to the basal layer via cell-cell adhesion to form the spinous cell layer. Moreover, compressive forces in cell monolayers of simple epithelia and of epidermal keratinocytes can promote extrusion and stratification, respectively [29, 30]. Since uncoupling the desmosome/IF connection resulted in mechanical behaviors that could be consistent with a reduction in compressive forces being experienced by cells expressing DPNTP, we next examined the effects of DPNTP on stratification in NHEKs using two models of keratinocyte stratification.

To begin, we examined the effects of uncoupling the desmosome/IF linkage on the ability of ephrin-induced colonies to stratify (Figure 2A). The lateral expansion of NHEKs is suppressed upon addition of ephrin peptide, inducing colonies of cells to stratify and form multiple layers [15]. This results in the piling of cells and thus the percentage of overlapping nuclei can be used as a proxy to quantify stratification events. In control ephrin-induced colonies, an average of 25% of the nuclei were overlapping, indicating at least 25% of cells had stratified by day 7 (Figure 2B). In contrast, expression of DPNTP resulted in an average of 9% overlapping nuclei at this timepoint (Figure 2B). Since there was no significant difference in total cell density (Figure 2B), these data suggest that uncoupling the desmosome/IF linkage impedes the stratification process.

**Figure 2.**
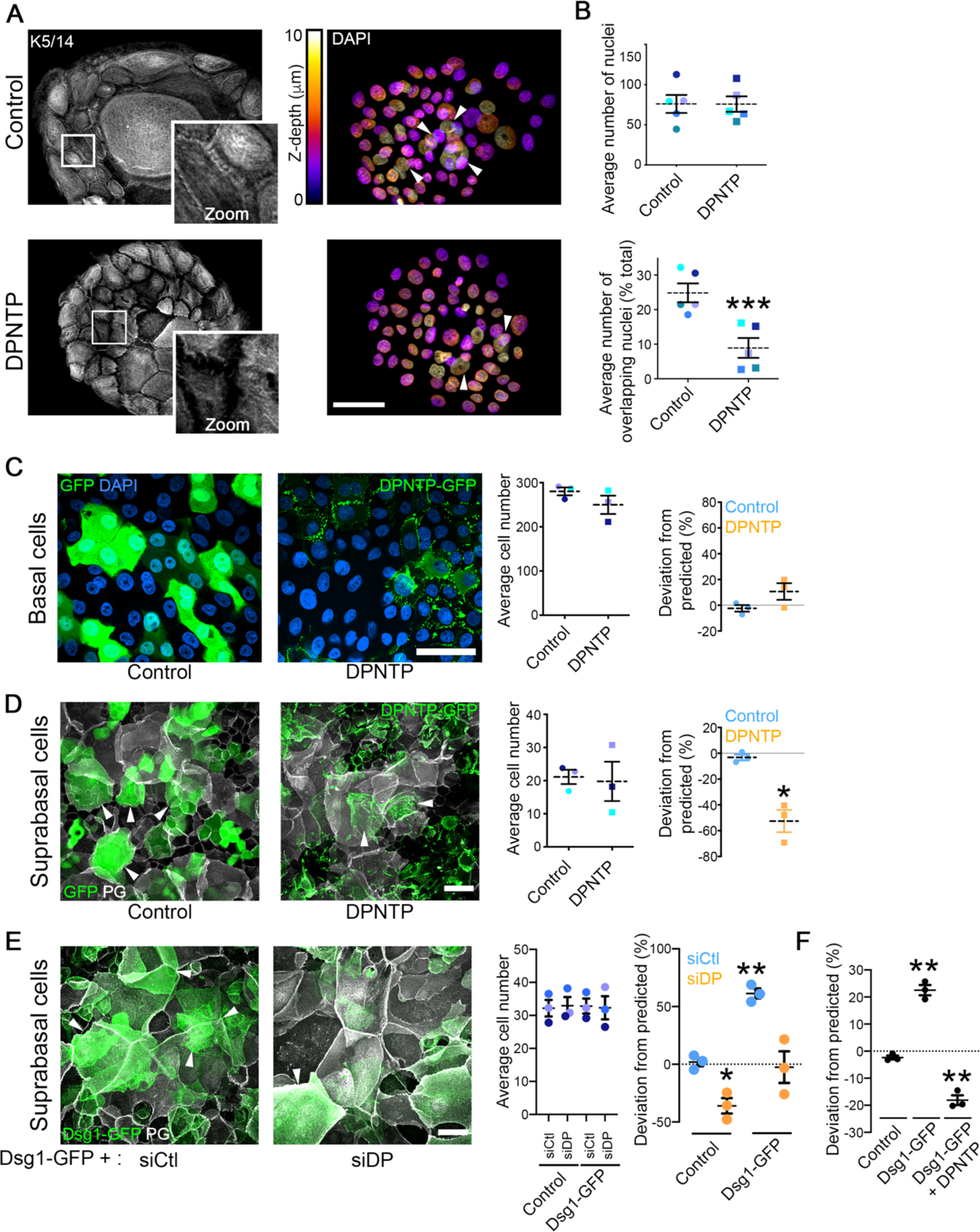
Uncoupling the desmosome/IF connection hinders epidermal keratinocyte stratification while promoting differentiation. A) Maximum projection micrographs show immunostaining of keratin 5/14 (K5/14) intermediate filaments and nuclei stained with DAPI using the indicated look-up table that represents z-depth in control and DPNTP-expressing ephrin colonies. Zooms show retraction of keratin filaments from cell-cell interfaces upon uncoupling the desmosome/intermediate filament with DPNTP. Bar is 50 μm. B) The average total number of DAPI-stained nuclei (as shown in A) per ephrin colony (top) and the average number of overlapping DAPI-stained nuclei (bottom) are shown for control and DPNTP-expressing ephrin colonies. Examples of overlapping nuclei are indicated with arrowheads in A. Dashed lines indicate the mean of 5 independent experiments and error bars are SEM. *p<0.0001, paired t-test. C) Unlabeled wild type NHEKs were mixed at known ratios with NHEKs expressing either GFP as a control or DPNTP-GFP and cultured in 1.2 mM calcium medium for 3 days. Left, representative maximum projection micrographs of the basal layer of cultures are shown using DAPI shown in blue to label the total cell population. Bar is 50 μm. Middle, quantification of the average cell number per field from 3 independent experiments is shown. Right, quantification of the deviation from the predicted representation of GFP-positive cells in the basal layer is shown. Dashed lines indicate the mean of 3 independent experiments and error bars are SEM. D) Unlabeled wild type NHEKs were mixed at known ratios with NHEKs expressing either GFP as a control or DPNTP-GFP and cultured in 1.2 mM calcium medium for 3 days. Left, representative maximum projection micrographs of the suprabasal layers of cultures are shown using plakoglobin (PG) to label the total cell population. Examples of GFP-positive suprabasal cells are indicated with arrowheads. Bar is 50 μm. Middle, quantification of the average cell number per field from 3 independent experiments is shown. Right, quantification of the deviation from the predicted representation of GFP-positive cells in the suprabasal layers is shown. Dashed lines indicate the mean of 3 independent experiments and error bars are SEM. *p=0.026, one sample t test with theoretical mean of 0. E) Unlabeled wild type NHEKs were mixed at known ratios with NHEKs expressing Dsg1-GFP treated with siCtl or siDP and cultured in 1.2 mM calcium medium for 1 day. Left, representative maximum projection micrographs of the suprabasal layer of cultures are shown using plakoglobin (PG) to label the total cell population. Examples of Dsg1-GFP positive suprabasal cells are indicated with arrowheads. Bar is 50 μm. Middle, quantification of the average cell number per field from 3 independent experiments is shown. Right, quantification of the average deviation from the predicted representation of Dsg1-GFP positive cells treated with the indicated siRNA in the suprabasal layer is shown for 3 independent experiments and error bars are SEM. *p=0.038 and **p=0.005, one sample t test with theoretical mean of 0. F) Unlabeled wild type NHEKs were mixed at known ratios with NHEKs expressing GFP (Control), Dsg1-GFP, and Dsg1-GFP co-expressing DPNTP-FLAG (DPNTP) and cultured in 1.2 mM calcium medium for 1 day. Quantification of the average deviation from the predicted representation in the suprabasal layer is shown for 3 independent experiments and error bars are SEM. **p*≤*0.001, one sample t test with theoretical mean of 0.

To better understand the function of the desmosome/IF linkage during stratification, we modeled the process using an in vitro competition assay. Fluorescently-labeled, genetically manipulated cells were mixed at known ratios with wild type cells to form a confluent monolayer that was then induced to stratify with a calcium switch (Figure S2A). Two days after calcium switching, the fate of the labeled cells was tracked. The percentage of labeled cells in the basal and suprabasal layers was compared to the percentage of cells initially plated for the assay (Figure S2B). If the genetic manipulation had no effect on the ability to compete with wild type cells to stratify, then there would be no deviation from the predicted percentage either in the basal or suprabasal layers. This was observed in control experiments where expression of cytoplasmic GFP had no significant effect on the calculated percentage of labeled cells in either the basal or suprabasal layers (Figure 2C and D). However, quantification of cells expressing the uncoupling mutant suggested a trend toward being retained in the basal layer and a significant decrease in the number of labeled cells in the suprabasal layer compared to what would be predicted (Figure 2C and D). There were no significant differences in cell density of basal or suprabasal cells in control and DPNTP-expressing conditions (Figure 2C and D). Additionally, depletion of endogenous DP using two distinct siRNA treatments resulted in a significantly decreased representation of labeled suprabasal cells while a non-targeting siRNA treatment had no significant effect (Figure S3A and B), corroborating the results obtained using the uncoupling mutant. Together, these data support a role for the desmosome/IF linkage during the stratification process such that breaking the connection renders epidermal keratinocytes less able to compete with wild type cells for sorting into the suprabasal layers.

We previously showed a role for Dsg1, a desmosomal cadherin expressed exclusively in stratified epithelia, in promoting delamination in 3D epidermal equivalents. Importantly, Dsg1 is sufficient to drive simple epithelial cells (MDCK) to exit a monolayer to form a second layer [31]. To determine if Dsg1-dependent movement into the suprabasal layers requires an intact IF connection, we ectopically expressed Dsg1-GFP in NHEKs and tracked their fate in the competition assay. Retroviral transduction of Dsg1-GFP into keratinocytes promoted a significant enrichment in the suprabasal layer, while GFP alone had no effect (Figure 2E). Moreover, the suprabasal enrichment of Dsg1-expressing cells was abolished upon either expression of DPNTP or siRNA-mediated DP loss (Figure 2E and F). Therefore, our data indicate that the desmosome/IF linkage is necessary for both basal compressive forces and for Dsg1-mediated sorting into the suprabasal layers. These data support the idea that our previously observed Dsg1-induced changes in cortical actin remodeling and reduced membrane tension require a DP-IF “clutch” to promote the efficient movement of newly differentiating keratinocytes expressing Dsg1 into the next layer.

### Uncoupling the desmosome/IF linkage promotes NHEK differentiation through the mechanosensitive SRF pathway

Generally, it is thought that the process of epidermal stratification and the program of terminal differentiation occur concurrently [32]. Therefore, we assessed the effects of uncoupling the desmosome/IF linkage on biochemical markers of differentiation in both 2D high calcium- and ephrin-induced models. Surprisingly, expression of DPNTP in both models resulted in a significant increase in the protein expression levels of differentiation markers including the desmosomal cadherins Dsg1 and desmocollin 1 (Dsc1), the suprabasal cell IF protein keratin 1, and the cornified envelope component loricrin (Figure 3A and B). There were no significant effects detected on the expression of total DP or on the classical cadherin E-cadherin, which are expressed in both basal and suprabasal cells. These data suggest that disengaging the desmosome/IF connection impedes stratification while accelerating differentiation.

**Figure 3.**
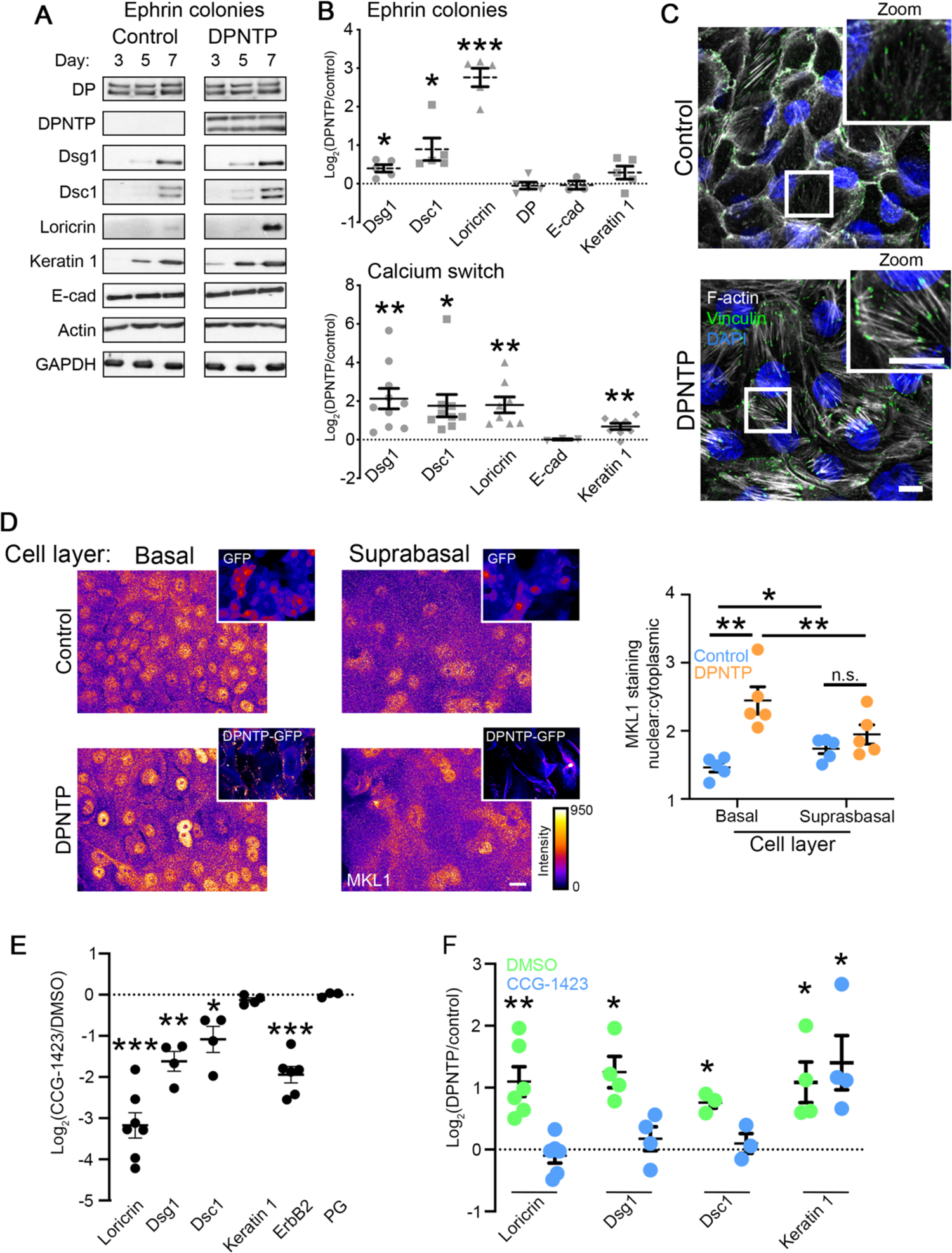
Uncoupling the desmosome/IF linkage promotes NHEK differentiation through the mechanosensitive SRF pathway. A) Western blots showing the expression of the indicated proteins during a differentiation time course of control or DPNTP-GFP expressing ephrin colonies. B) Upper, quantification of the fold change (Log_2_-transformed) of DPNTP-expressing over control ephrin colonies at day 7 is shown for expression of the indicated proteins. Means are from 3-5 independent experiments and error bars are SEM. *p*≤*0.04, ***p=0.0003, one sample t test with theoretical mean of 0. Lower, quantification of the fold change (Log_2_-transformed) of DPNTP-expressing over control monolayer cultures exposed to 1.2 mM calcium medium for 3-4 days are shown for indicated protein expression. Means are from 3-10 independent experiments and error bars are SEM. *p=0.016, **p*≤*0.004, one sample t test with theoretical mean of 0. C) Maximum projection micrographs show F-actin (using phalloidin) and immunofluorescence staining of vinculin near the basal/substrate interface of differentiating monolayers of NHEKs expressing either GFP (Control) or DPNTP-GFP at 5 days after calcium switch. DAPI indicates nuclei in blue and bar is 10 μm. D) Left, maximum projection micrographs show MKL1 immunofluorescence staining in the basal and suprabasal layers of day 5 cultures of NHEKs using the indicated lookup table. Bar is 10 μm. Insets show GFP and DPNTP-GFP expression. Right, quantification of the nuclear to cytoplasmic ratio of MKL1 staining is shown for cultures expressing GFP (Control) or DPNTP-GFP in the indicated cell layers. *p=0.021, **p*≤*0.009, paired t-test. E) Quantification of the fold change (Log_2_-transformed) of cultures treated with the SRF inhibitor CCG-1423 over DMSO-treated controls at day 5 is shown for the indicated proteins. Means are from 3-7 independent experiments and error bars are SEM. *p=0.0423, **p=0.0065, ***p*≤*0.0002, one sample t test with theoretical mean of 0. F) Quantification of the fold change (Log_2_-transformed) of DPNTP-expressing over GFP control keratinocytes is shown for the indicated protein expression for both DMSO and CCG-1423 treatment. Means are from 3-6 independent experiments and error bars are SEM. *p*≤*0.05, **p=0.006, one sample t test with theoretical mean of 0.

Towards understanding how differentiation is accelerated in DPNTP expressing keratinocytes, we explored potential mechanosensitive pathways known to regulate epidermal differentiation. One such pathway centers on SRF (serum response factor), which is regulated by the ratio of G- and F-actin. MKL1 (also known as MAL) is a transcriptional coregulator of SRF that normally binds to G-actin and is retained in the cytoplasm. With increased F-actin content, G-actin is released from MKL1, which in turn redistributes into the nucleus where it can work with SRF to induce keratinocyte differentiation [33]. Moreover, we previously showed that SRF is upstream of Dsg1 in this pathway [34]. Therefore, we considered the role of SRF in the DPNTP-mediated acceleration of differentiation.

To address this possibility, we uncoupled the desmosome/IF linkage in NHEK monolayers with DPNTP. This uncoupling resulted in decreased cortical F-actin staining with a concomitant increase in observed focal adhesion-anchored F-actin stress fibers at the basal surface (Figure 3C). These results are consistent with the observed role of DP in the regulation of the F-actin cytoskeleton in mouse studies [35–37]. To determine if SRF signaling itself was affected downstream of desmosome/IF modulation, we assessed MKL1 nuclear localization. In the limited number of DPNTP-expressing stratified cells in these cultures, we did not see differences in the nuclear/cytoplasmic ratio of MKL1 staining compared to stratified controls (Figure 3D). However, DPNTP expression led to an enrichment of MKL1 in the nuclei of basal cells (Figure 3D), suggesting a precocious activation of the SRF pathway. Consistent with this idea, CCG-1423, a small molecule inhibitor of SRF, reduced the expression of Dsg1, Dsc1, and loricrin as well as abolished the DPNTP-mediated increase in expression of these proteins (Figure 3E and F).

### The DPNTP-mediated effects on differentiation are dependent on ErbB2

Our results above showed that uncoupling the desmosome/IF linkage disconnects the coordination of the formation of a second cell layer and induction of differentiation through the SRF pathway, both early morphogenic events linked to Dsg1 dependence. Dsg1 is expressed first as basal cells commit to differentiate and its expression progressively increases, so that it is most concentrated in the superficial layers of the epidermis. This led us to assess potential transitional signaling platforms.

Based on our previous findings showing that Dsg1 tunes EGFR signaling early in differentiation, we first considered the ErbB family of receptor tyrosine kinases. There are four members of the ErbB family (ErbB1-4) and all have been reported to be expressed in human skin [38–40]. EGFR (i.e. ErbB1) is highly active in the basal layer where it keeps cells in an undifferentiated state [41]. Dsg1 promotes the transition of the undifferentiated basal phenotype to a differentiated suprabasal one through modulation of EGFR/Erk activity [2, 41] and promotion of delamination [31]. Expression levels of ErbB2 increase as keratinocytes differentiate [38], but its role in this process is not known. To address the potential role of ErbB2 in differentiation, we utilized a pharmacological approach in which we treated cells with the ErbB2 specific inhibitor TAK165 [42]. TAK165 treatment significantly reduced both total and Y877 phosphorylated ErbB2 (Figure S4A), a site within the kinase domain that has been reported to increase ErbB2 activity [43, 44]. Addition of TAK165 also resulted in a significant decrease in the protein expression levels of the differentiation associated proteins Dsg1, Dsc1, and loricrin, while no effects were detected for the classical cadherin, E-cadherin (Figure S4A). These data suggest a potential feedback loop between ErbB2 activity and desmosomal protein expression. Moreover, uncoupling the desmosome/IF linkage promoted an increase in phosphorylation of Y877 ErbB2 (Figure S4B) and this increase was abrogated upon depletion of Dsg1 (Figure S4C). Treatment of NHEKs with TAK165 abolished the differences seen in differentiation between DPNTP and controls treated with DMSO (Figure S4D and E), suggesting that the DPNTP-mediated increase in expression of differentiation related proteins is dependent on ErbB2. These data indicate that ErbB2 plays a role in the differentiation process of NHEKs, consistent with a previously reported decrease in keratin 10 and filaggrin in ErbB2-deficient mouse epidermis [45]. Moreover, we have identified ErbB2 as a putative new target of SRF signaling, as treatment of wild type NHEKs with CCG-1423 results in a significant decrease in ErbB2 expression (Figure 3E).

### Desmosomes regulate a tension gradient and TJ proteins in epidermal equivalents

To address whether desmosomes regulate mechanical forces and cell behaviors in the upper layers, where TJs form, we assessed indicators of mechanical force throughout the suprabasal layers. For these experiments we took a more targeted approach, ablating the function of the desmosomal cadherin Dsg1, which is specifically expressed in the stratified layers of the epidermis. Vinculin is a tension sensitive cell-cell junction component recruited to actin-anchored adherens junctions [46]. In control 3D epidermal equivalent cultures infected with nontargeting shRNA, vinculin immunostaining was restricted to the SG layer where tension is thought to be high [2]. This idea is supported by laser ablation experiments of suprabasal cells in 3D epidermal equivalents, which unlike basal cells at day 6, expand upon ablation (Movie S1). Control cultures exhibited a steep gradient of vinculin border staining along the apical to basal axis, which was attenuated in Dsg1-deficient cultures, with increased cell-cell border localized vinculin in both the spinous and basal layers (Figure 4A). These data suggest that Dsg1 loss shifts the gradient of forces within the epidermal culture, similar to results obtained in E-cadherin-deficient mouse epidermis [2].

**Figure 4.**
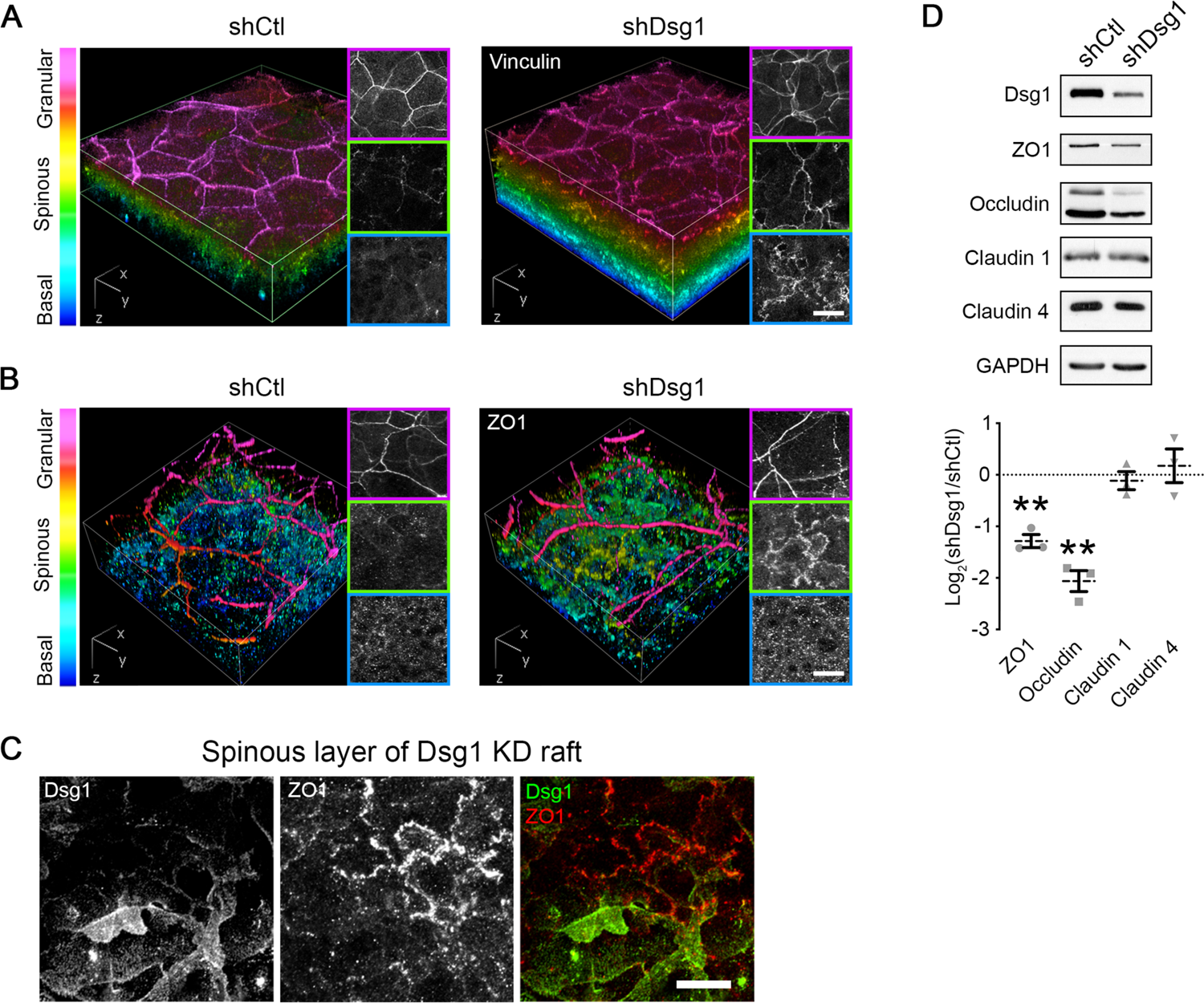
Dsg1 regulates a tension gradient and tight junction proteins in epidermal equivalents. A) 3D renderings of whole mount immunostaining of tension-sensitive vinculin in control (shCtl) and Dsg1-depleted (shDsg1) day 6 epidermal equivalent cultures are presented with a look-up table that depicts z-depth in the indicated colors. Right panels show representative vinculin staining per layer. Bar is 20 μm. B) 3D renderings of whole mount immunostaining of tight junction component ZO1 in control (shCtl) and Dsg1-depleted (shDsg1) day 6 epidermal equivalent cultures are presented with a look-up table that depicts z-depth in the indicated colors. Right panels show representative ZO1 staining per layer. Bar is 20 μm. C) Representative micrographs of spinous layer in Dsg1-depeleted day 6 epidermal equivalent cultures showing both ZO1 and Dsg1 immunostaining. Bar is 20 μm. D) Upper, western blot analysis from control (shCtl) and Dsg1-depleted (shDsg1) day 6 epidermal equivalent cultures show the protein expression of the indicated tight junction proteins with GAPDH shown as a loading control. Lower, quantification of the fold change (Log_2_-transformed) of Dsg1-depleted over control cultures are shown for the indicated proteins. Dashed lines indicate the mean of 3 independent experiments and error bars are SEM. **p<0.01, one sample t test with theoretical mean of 0.

Since TJs are thought to be regulated by a tension gradient in the epidermis [2], we next assessed the staining of the TJ component ZO1 in epidermal equivalent cultures. ZO1 immunolabeling in control cultures shows specific localization to the SG layer (Figure 4B), consistent with the reported location where functional epidermal TJs form [5]. Similar to the effects observed with vinculin redistribution, Dsg1 depletion led to a shift in this staining pattern (Figure 4B). A shift in the ZO1 staining pattern was also observed upon treatment with a Rho activator, whereby ZO1 similarly localized prematurely to cell-cell interfaces in the spinous layer (Figure S5A and B). The effect of Dsg1 on ZO1 appears to be cell autonomous, as mosaic expression of Dsg1 in the spinous layer altered cell-cell contact localized ZO1 specifically in Dsg1-depleted cells (Figure 4C). Moreover, biochemical analysis of the TJ proteins ZO1 and occludin indicates Dsg1 depletion results in a decrease in their expression (Figure 4D), while no effects were detected for claudin 1 or claudin 4.

### Uncoupling the desmosome/IF connection diminishes epidermal keratinocyte barrier function

In multiple models, TJs and their protein components have been suggested to be sensitive to mechanical input [2, 47–49]. We hypothesize the patterning of the desmosome/IF components drives polarized, layer-specific epidermal functions; thus we next assessed the effects of uncoupling the desmosome/IF linkage on the ensuing development of the epidermal TJ barrier. Staining of the TJ component occludin in sagittal sections of epidermal equivalent cultures indicated that expression of DPNTP resulted in a shift to membrane enrichment of occludin in the spinous layer compared with controls, while no differences were seen in the SG layer (Figure S6A). These results are consistent with previous work showing that loss of DP in mouse epidermis modulates TJ protein expression and localization [50]. Together with data from Dsg1-deficient cultures, these results suggest that the desmosome/IF connection controls the distribution of forces within the epidermis to restrict TJ protein localization to the SG layer.

Since uncoupling the desmosome/IF linkage affects TJ component localization and the mechanical properties of cells, we hypothesized this could result in alterations in the TJ barrier function, which is attributed to the outer layers of the epidermis (Figure 5A). To address this, we generated human epidermal equivalent cultures using a porous transwell system (Figure S1D). We then performed transepidermal electrical resistance (TEER) measurements on the developing cultures as a means of assessing barrier function (Figure 5B). We found that altering cell mechanics in this 3D system using an activator of actomyosin contractility (CN01, a Rho activator) had an effect on the organization of F-actin in all layers of the cultures and significantly inhibited barrier function (Figure S5C and D). In addition, we performed immunostaining of the TJ protein ZO1 as an indicator of the functional TJs found in the SG layer of the epidermis [5]. ZO1 localizes to cell-cell interfaces in the SG layer in control GFP-expressing cultures, an area with strong F-actin staining (Figure 5C). However, when desmosomes are uncoupled from IF with DPNTP, there is a cell autonomous loss of ZO1 staining in the SG layer (Figure 5D), suggesting a potentially impaired barrier. We do not observe altered ZO1 staining in the basal or spinous layers upon expression of DPNTP (Figure S6B), suggesting that layer-specific mechanisms exist. TEER measurements of the developing barrier were significantly reduced in DPNTP-expressing cultures compared with GFP controls (Figure 5E), indicating expression of DPNTP reduces epidermal barrier function.

**Figure 5.**
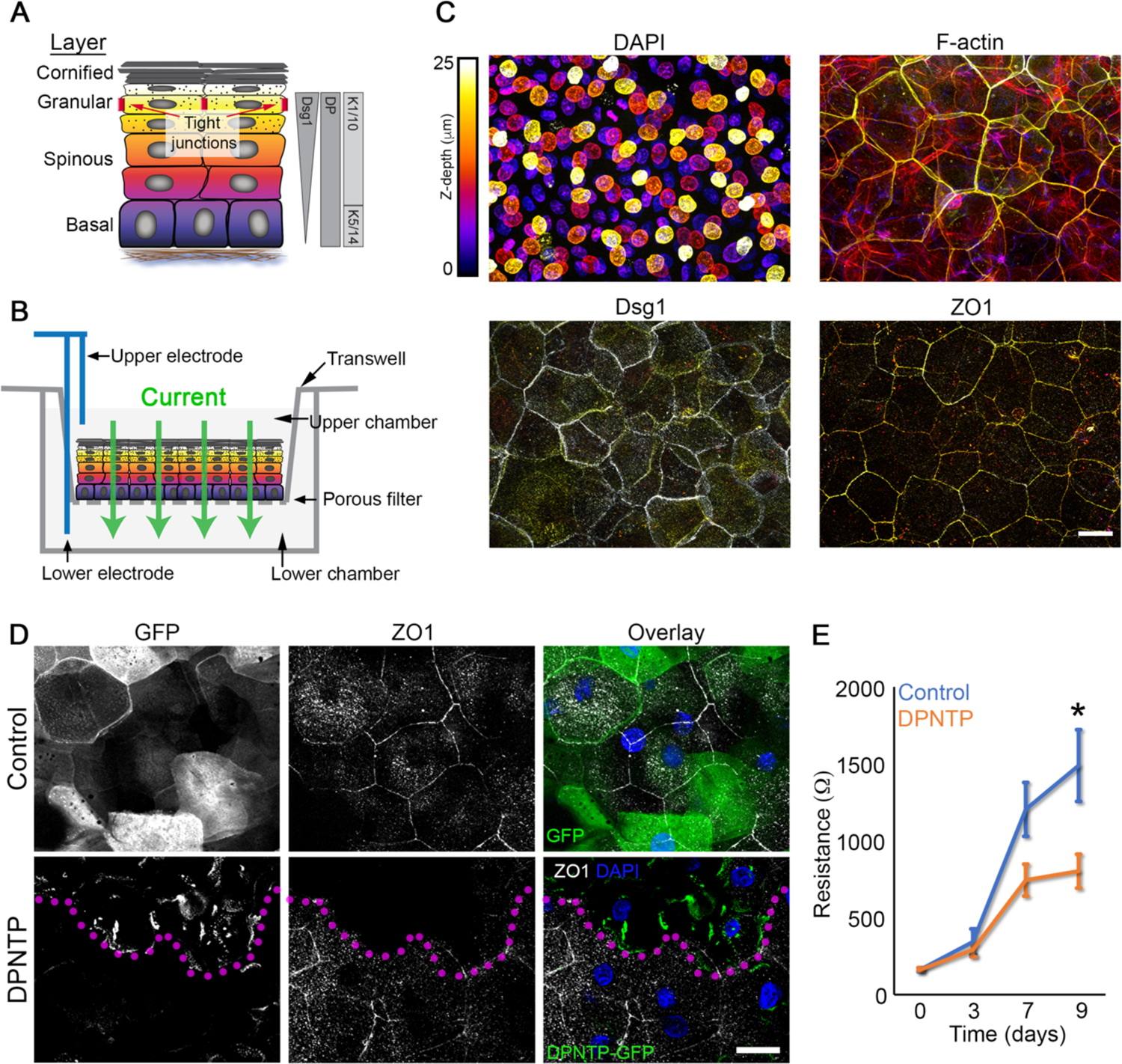
Uncoupling the desmosome/IF connection diminishes epidermal keratinocyte barrier function. A) Schematic shows the layers of the epidermis and functional tight junctions localize to the second granular layer. The expression patterns of desmoplakin (DP), keratins (K), and desmoglein 1 (Dsg1) are also shown. B) Schematic shows how transepidermal electrical resistance (TEER) experiments were performed on transwell epidermal equivalent cultures. Electrodes were used to pass a current through the 3D culture that was grown on a transwell insert and resistance measurements were taken as an indicator of barrier function. C) Maximum projection micrographs of day 9 transwell epidermal equivalent cultures show staining for nuclei (DAPI), F-actin (phalloidin), and immunostaining for the indicated proteins using a look-up table indicating z-depth. Bar is 25 μm. D) Fluorescence micrographs show the expression of GFP (Control) and DPNTP-GFP in the granular layer of day 9 transwell epidermal equivalent cultures that were also immunostained for the tight junction component ZO1. Magenta dotted line delineates DPNTP-GFP high expressing cells. Nuclei are stained with DAPI in blue. Bar is 25 μm. E) Quantification of resistance measurements from TEER experiments performed on a time course of control (GFP) and DPNTP-GFP expressing transwell epidermal equivalent cultures. *p=0.0125, Sidak’s multiple comparison test from 7 independent experiments. Data are presented as mean ± SEM.

### ErbB2 as a potential mediator of desmosome-mediated effects on epidermal polarity and barrier function

Having established that ErbB2 mediates differentiation downstream of SRF signaling, we considered the possibility that it also contributes to later events that drive barrier function. Toward this end, we more closely assessed the relative expression and localization of EGFR and ErbB2 in human epidermis. ErbB2 localization was enriched in the SG2 layer of human epidermis (Figure 6A), the specific layer where functional TJs form [5]. While total ErbB2 was expressed at all timepoints evaluated, its phosphorylation status at Y877 increased as human epidermal equivalent cultures developed (Figure 6B). Moreover, ErbB2 has been reported to modulate epithelial permeability, being linked to both loss and gain of barrier function [51], indicative of context-dependent functions. This suggested to us that ErbB2 may play a role in the desmosome-mediated regulation of TJs within the epidermis.

**Figure 6.**
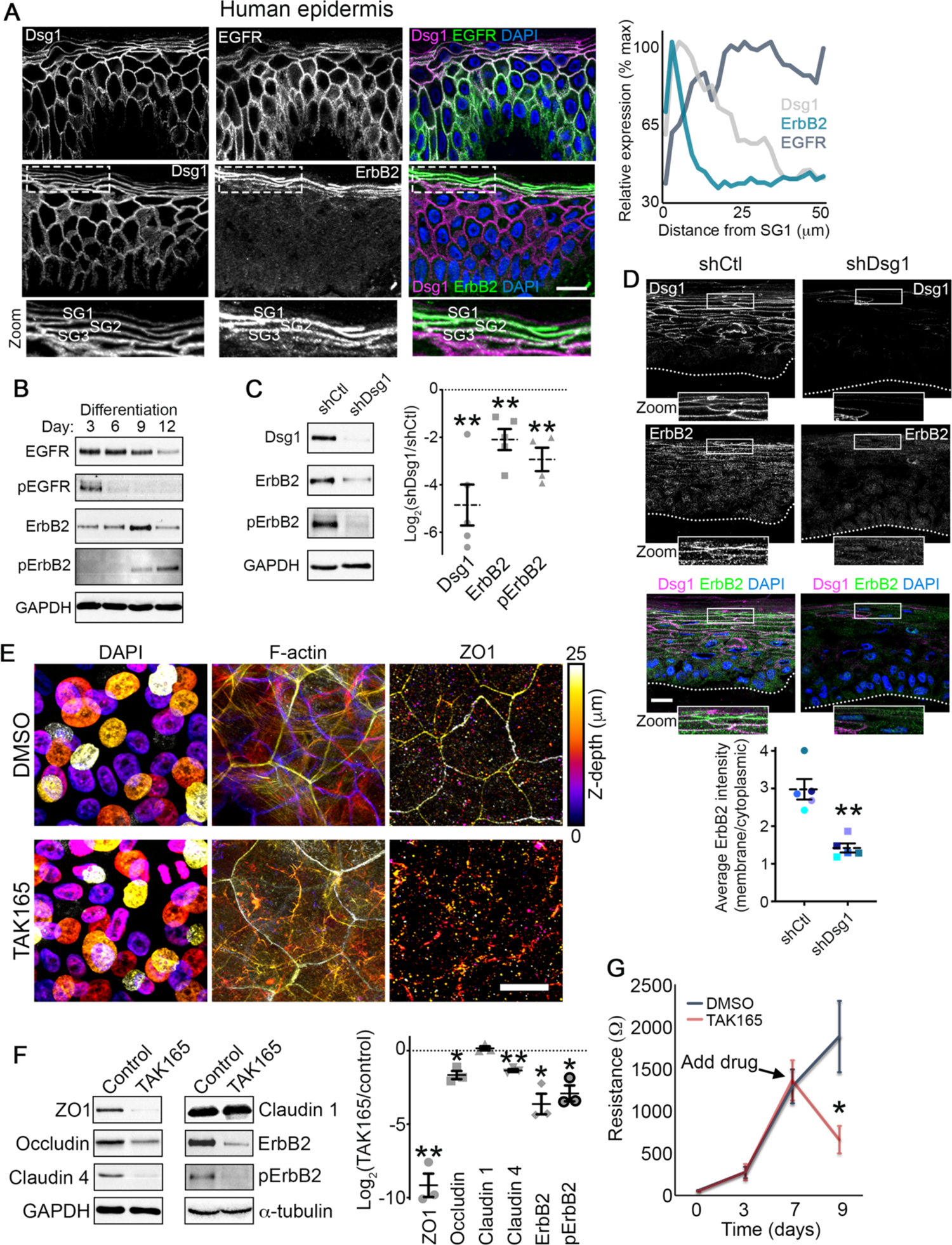
ErbB2 as a potential mediator of Dsg1-mediated effects on epidermal polarity and barrier function. A) Left, representative micrographs of transverse sections of human epidermis immunostained for the indicated proteins are shown. Zoom is of area in dashed box highlighting the stratum granulosum (SG1-3). DAPI-stained nuclei are in blue and bar is 20 μm. Right, line scan analysis shows the average relative level of fluorescence intensity from 3 independent samples for the indicated proteins as a function of distance from the most superficial living epidermal layer (stratum granulosum, SG). B) Representative western blots of a differentiation time course of human epidermal equivalent cultures showing the total EGFR and Y1068-phosphorylated EGFR (pEGFR) as well as total ErbB2 and Y877-phosphorylated ErbB2 (pErbB2). GAPDH is used as a loading control. C) Left, representative western blots of day 6 human epidermal equivalent cultures expressing either a non-targeting shRNA (shCtl) or an shRNA targeting Dsg1 (shDsg1) showing Dsg1 protein level, total ErbB2, and Y877-phosphorylated ErbB2 (pErbB2). GAPDH is used as a loading control. Right, quantification of the fold change (Log_2_-transformed) of Dsg1-depleted over control cultures are shown for the indicated proteins. Dashed lines indicate the mean of at least 4 independent experiments and error bars are SEM. **p<0.01, one sample t test with theoretical mean of 0. D) Micrographs of transverse sections of day 6 human epidermal equivalent cultures expressing either a non-targeting shRNA (shCtl) or an shRNA targeting Dsg1 (shDsg1) immunostained for Dsg1 and ErbB2 are shown. Dotted line marks the bottom of the basal layer. DAPI-stained nuclei are in blue and bar is 20 μm. Lower, quantification of ErbB2 immunofluorescence intensity expressed as a ratio of the membrane localized signal over the cytoplasmic localized signal. Dashed lines indicate the mean of 5 independent experiments and error bars are SEM. **p=0.008, paired t test. E) Day 9 transwell epidermal equivalent cultures were treated with either DMSO or the ErbB2 inhibitor TAK165 (1 μM) for the last 48 hours prior to harvesting. Maximum projection micrographs show staining for nuclei (DAPI), F-actin (phalloidin), and immunostaining for the tight junction protein ZO1 using a look-up table that indicates z-depth. Bar is 25 μm. F) Day 9 transwell epidermal equivalent cultures were treated with either DMSO or the ErbB2 inhibitor TAK165 (1 μM) for the last 48 hours prior to harvesting. Left, western blots showing the expression of the indicated tight junction proteins for samples treated with either DMSO (Control) or TAK165. GAPDH and *α*-tubulin are used as loading controls. Right, quantification of the fold change (Log2-transformed) of TAK165-treated over control cultures are shown for the indicated proteins. Dashed lines indicate the mean of 3 independent experiments and error bars are SEM. *p=0.025, **p<0.01, one sample t test with theoretical mean of 0. G) Quantification of resistance measurements from TEER experiments performed on a time course of transwell epidermal equivalent cultures that were treated with either DMSO or the ErbB2 inhibitor TAK165 (1 μM) on day 7 (as indicated) for the last 48 hours. *p=0.0325, paired t test from 5 independent experiments. Data are presented as mean ± SEM.

To examine the effects of Dsg1 on ErbB2 function, we performed Dsg1 depletion experiments. Loss of Dsg1 resulted in a biochemical decrease in both total and Y877 phosphorylated ErbB2 as well as a loss of ErbB2 at sites of cell-cell contact in the upper layers of human epidermal equivalent cultures (Figure 6C and D). These data suggest that Dsg1 indeed regulates ErbB2. We then assessed the potential role of ErbB2 in regulating TJ protein expression and function. Since we showed that ErbB2 is important for the process of epidermal keratinocyte differentiation (Figure S4A), we allowed human transwell epidermal equivalent cultures to form for 7 days and then treated with either DMSO as a control or the TAK165 inhibitor for 2 additional days. TAK165 treatment altered overall ZO1 and F-actin staining (Figure 6E) with aberrant punctate ZO1 staining in the spinous layer (Figure S7), similar to the results seen upon Dsg1 depletion (Figure 4B and C). Additionally, compared to controls, TAK165 significantly reduced the expression of TJ protein components including ZO1, claudin 4, and occludin (Figure 6F). Finally, inhibiting ErbB2 with TAK165 significantly diminished the forming barrier assessed using TEER measurements (Figure 6G).

ErbB2 has been reported to be phosphorylated on Y877 by Src family kinases (SFKs), which increases its kinase activity [44]. In human epidermis, SFKs are expressed in patterns with Src expression largely restricted to the basal layer while Fyn and Yes are expressed in the granular layer (Figure 7A). This suggested to us that SFKs may be responsible for desmosome-mediated alterations in Y877 phosphorylated ErbB2. Indeed, treatment of day 7 epidermal equivalents with the SFK inhibitor PP2 resulted in a significant decrease in Y877 p-ErbB2 (Figure 7B). Moreover, in Dsg1 gain-of-function experiments, retroviral transduction of ectopic, FLAG-tagged Dsg1 into NHEKs resulted in a significant increase in the ratio of Y877 p-ErbB2 to total ErbB2 (Figure 7C). This gain-of-function as well as the DPNTP-mediated increase in p-ErbB2 are lost upon inhibition of SFKs with PP2 (Figures 7C and D). However, Dsg1 overexpression is still able to promote an increase in the relative amount of Y877 p-ErbB2 upon SRF inhibition suggesting Dsg1 is downstream of SRF and upstream of ErbB2 (Figure 7C).

**Figure 7.**
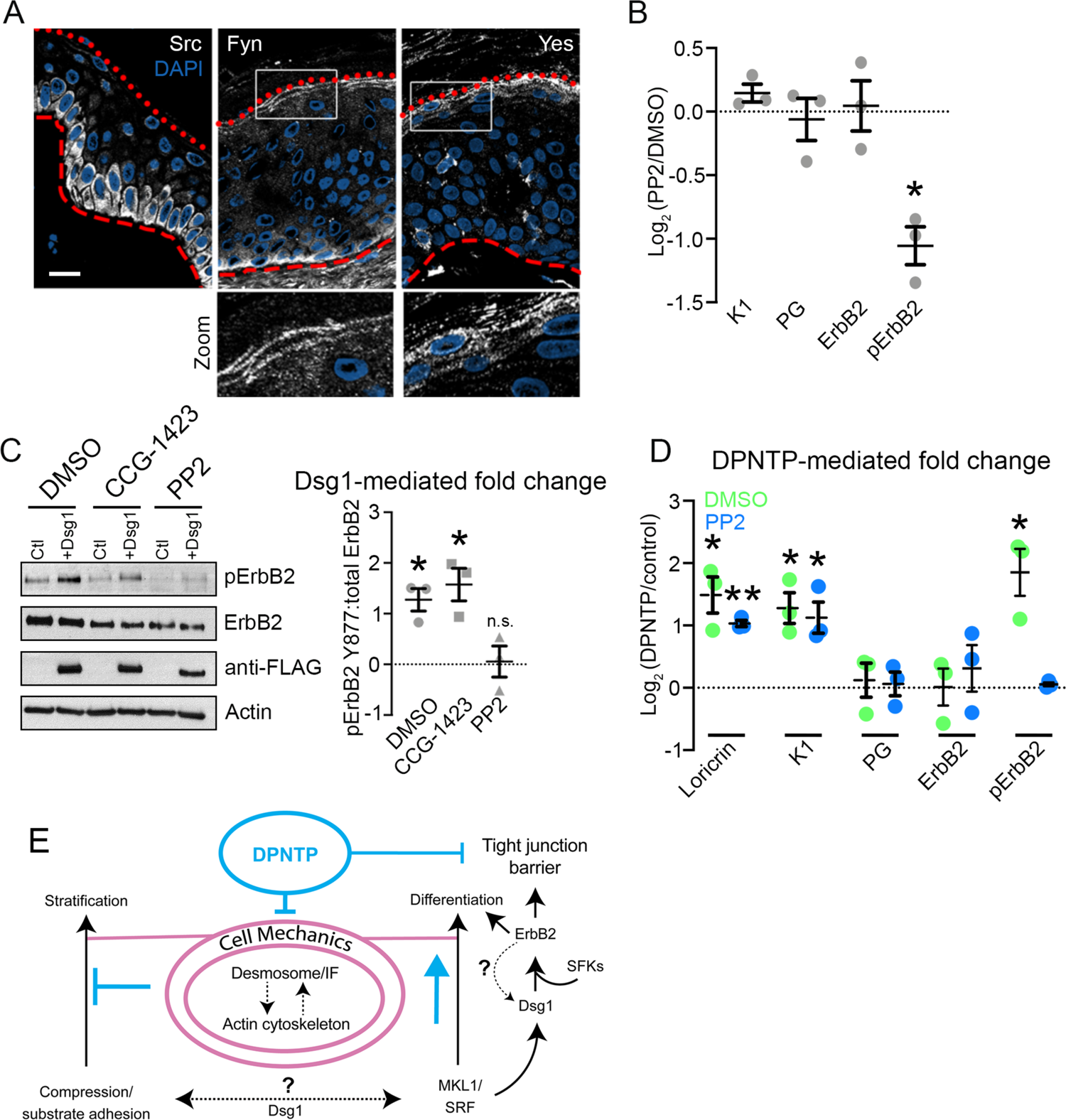
Desmosome-mediated modulation of ErbB2 phosphorylation depends on Src family kinases. A) Representative micrographs of transverse sections of human epidermis immunostained for the indicated Src family kinases (SFKs) are shown. Red dotted line marks the top of the granular layer and the dashed red line denotes the bottom of the basal layer. DAPI-stained nuclei are in blue and bar is 20 μm. B) Quantification of the fold change (Log_2_-transformed) of cultures treated with the SFK inhibitor PP2 (6 hours prior to harvest) over DMSO-treated controls at day 5 are shown for the indicated proteins. Means are from 3 independent experiments and error bars are SEM. *p=0.0195, one sample t test with theoretical mean of 0. C) Left, western blots showing the expression of the indicated proteins in day 5 cultures of NHEKs expressing either GFP (Ctl) or Dsg1-GFP and treated with either the SRF inhibitor CCG-1423, the SFK inhibitor PP2, or DMSO as a control. Right, quantification of the fold change (Log_2_-transformed) in the ratio of pErbB2:total ErbB2 is shown for the indicated treatments. Means are from 3 independent experiments and error bars are SEM. *p*≤*0.04, one sample t test with theoretical mean of 0. D) Quantification of the fold change (Log_2_-transformed) of DPNTP-expressing over GFP control keratinocytes are shown for the indicated protein expression for both DMSO and PP2 treatment. Means are from 3 independent experiments and error bars are SEM. *p*≤*0.05, **p=0.0023, one sample t test with theoretical mean of 0. E) Working model of the effects of the desmosome/intermediate filament (IF) network on early and late epidermal morphogenetic events. During early morphogenesis, the desmosome/IF network works in conjunction with the actin cytoskeleton to regulate stratification through modulating the mechanical properties of basal cells. Uncoupling the desmosome/IF linkage with DPTNP reduced compressive forces and increased substrate adhesion, hindering stratification. Differentiation of keratinocytes proceeds through the mechanosensitive SRF pathway, which is accelerated when desmosomes are uncoupled from IF. This pathway includes Dsg1, which through SFKs modulates ErbB2 phosphorylation. ErbB2 in turn is critical to the formation of the epidermal barrier.

## Discussion

Apicobasal polarity is an intrinsic property of epithelial tissues that permits discrimination between interior and exterior compartments by establishing a functional barrier near the apical, external-facing surface. In contrast to simple epithelia, in which single cells in an epithelial sheet are polarized in an apicobasal fashion, in the epidermis apicobasal polarity is organized across multiple layers [10]. Here, we demonstrate that the desmosome/IF network is a critical player in establishing the polarized structure and function of the epidermis. Alterations in both basal and superficial cell mechanics in response to desmosome impairment are consistent with a switch from strong cell-cell forces to enhanced cell-substrate forces (basal layer) or enhanced intercellular tension in the cells below the TJ layer. For instance, in small colonies of differentiating keratinocytes the desmosome/IF DP uncoupling mutant resulted in an increase in vinculin-positive, cell-substrate contacts and associated F-actin stress fibers (Figures 1C and 3C), previously documented to signal a switch from strong cell-cell forces to enhanced cell-substrate forces [17–19]. At the same time, a decrease in cell-cell forces is supported by laser ablation experiments of early stage epidermal cultures (Figure 1D-F) and by our previous finding that expression of this mutant in A431 cells reduced cell-cell tugging forces [12].

How does interfering with basal cell mechanics affect an epidermal keratinocyte’s commitment to differentiate? During development of the epidermis, cell proliferation is thought to induce crowding and thus compressive forces that have been linked to epidermal fate specification and delamination [52–55]. However during homeostasis of adult mouse epidermis, inhibiting cell division, and thus reducing cell density, did not affect the ability of keratinocytes to delaminate [56]. Therefore, compressive forces in the basal layer appear to have different roles depending on the stage of development. In vitro, differentiating keratinocytes exhibit a transient decrease in cortical tension that is associated with the onset of expression of differentiation markers and subsequent exit of cells from the basal layer via delamination [57]. Here, we show that as normal human epidermal cultures progress from early to later stages of development, cells transition from a high-tension state to a compressive state (Figure 1D). These states are not achieved when IF are uncoupled from desmosomes. Thus, uncoupling IF from the desmosome reduces compressive forces within the basal layer that have been associated with differentiation. Consistent with this, cells were less capable of stratification into the superficial layers when IF were uncoupled from desmosomes (Figure 2D). These findings are supported by a recent publication that established a role for desmosomes and their attachment to IF during cell extrusion in a simple epithelial model [58]. This actomyosin-dependent process is often compared with stratification of multilayer epithelia [58]. Altogether, these data are consistent with a role for desmosomes and IFs working cooperatively with actomyosin to modulate the ability of a cell to leave the basal layer of an epithelium.

Dsg1 redistributed molecular tension in keratinocyte monolayers through rearrangements of cortical F-actin mediated by Arp2/3-dependent actin polymerization [31]. In the context of early morphogenesis, these tension-redistributing forces help promote basal cell delamination by reducing tension on E-cadherin [31]. Moreover, our previous observations show that the actomyosin contractile system is required for the altered mechanics observed when the desmosome/IF connection is impaired [12]. Furthermore, either uncoupling desmosomes from IF or siRNA-mediated depletion of DP abrogates the ability of Dsg1-positive cells to become enriched in the suprabasal layer (Figure 2E and F). Collectively, these data support the idea that desmosomes cooperate with the F-actin cytoskeleton to coordinate changes in cortical tension and adhesion necessary to drive stratification.

The process of keratinocyte stratification is generally associated with the onset of a terminal program of differentiation. Therefore, it was initially surprising that while uncoupling IF from desmosomes impedes the movement of keratinocytes into the suprabasal layers, the biochemical program of differentiation is accelerated (Figure 3A and B). This observation is not without precedent, as inhibition of actomyosin contractility was shown to uncouple morphogenesis and fate specification of inner cell mass cells in the developing mouse embryo [59]. We propose that uncoupling the desmosome/IF linkages precociously activates the SRF pathway to accelerate the process of differentiation, but due to a loss of “clutch” activity, altered cell-cell adhesion, and increased cell-substrate adhesion, they are less likely to stratify (Figure 7E).

Junctional tension, like that produced by actomyosin forces, recruits the TJ components ZO1 and occludin to promote barrier function in simple polarized epithelia [47, 49]. Dsg1 loss perturbed the normally restricted pattern of vinculin in the high-tension layer where TJs form, consistent with the idea that higher tension cell-cell junctions occurred in the lower layers (Figure 4A). This is in line with previous work showing impaired desmosome function results in increased molecular indicators of tension experienced by adherens junctions in mouse epidermis [50]. Conversely, multiple reports using E-cadherin knockout mouse models have shown that loss of epidermal E-cadherin does not have major effects on overall desmosome structure nor on markers of differentiation [60, 61]; however, combined loss of both epidermal E- and P-cadherin does [57]. In addition, interfering with Dsg1 shifts the localization ZO1 to more basal layers (Figure 4B and C). Coupled with previous work showing the importance of E-cadherin in the TJ barrier [2], these data are consistent with a model in which loss of Dsg1 induces a transfer of forces experienced to non-desmosomal junctions and premature mechanosensitive recruitment of junctional proteins to cell-cell interfaces in spinous layers. Moreover, disruption of the desmosome/IF linkage, which decreases cell-cell forces, is sufficient to reduce barrier function of transwell epidermal equivalent cultures (Figure 5E). Therefore, the fine tuning of mechanical input is critical to TJ barrier function and this tuning requires desmosome/IF attachment. Finally, perturbations of TJ proteins including claudins and occludin in epidermis affects keratinocyte differentiation [62–64], suggesting potential feedback loops to the regulation of differentiation-dependent desmosomal proteins.

ErbB2 activity is sensitive to changes in cell mechanics [41, 65, 66] and our data provide evidence suggesting that ErbB2 is a novel target of SRF signaling (Figure 3E), placing it downstream of this mechanosensitive pathway. In the epidermis, expression of ErbB2 is enriched in the superficial layers of the epidermis, where Dsg1 is at peak expression (Figure 6A). Depletion of Dsg1 abrogates the membrane localization of ErbB2 and the levels of Y877 phosphorylated ErbB2 (Figure 6B-D). We show that expression of either DPNTP or ectopic Dsg1 is sufficient to increase the relative amount of Y877 phosphorylated ErbB2 (Figures 7C-D and S4B). This site has been previously reported to be phosphorylated by SFKs [44] and here we show that both the DPNTP- and Dsg1-mediated increase in p-ErbB2 is abolished upon SFK inhibition with PP2 (Figure 7C-D). Since SFKs are expressed in patterns in the epidermis (Figure 7A), this suggests that layer-specific functions of the desmosome/IF network could be facilitated through modulation of restricted SFK activity.

Our previous work demonstrated that the onset of differentiation stimulated by Dsg1 expression in basal cells depends on its ability to dampen EGFR/Erk signaling that is active in the basal layer, which required an interaction between Dsg1 and Erbin (ErbB2 interacting protein) [41, 65]. Thus, together with the data presented here, this suggests a model in which a Dsg1 expression gradient functions to dampen EGFR basally to promote differentiation while stimulating ErbB2 suprabasally to support a functional TJ barrier. However, it is possible that ErbB2 coordinates with other members of the ErbB family to regulate TJ function and the role of Erbin in this process remains an avenue for future studies.

Collectively, the data presented here expose the desmosome/IF network as an essential element of the machinery in epidermis that cooperates with the F-actin cytoskeleton to pattern layer-specific mechanical properties. In the basal layer, this network affects compressive forces that coordinate the processes of stratification and differentiation; in the upper layer, the network integrates mechanosensitive cytoskeletal and signaling components to maintain the TJ barrier. We identify ErbB2 as a critical regulator of both keratinocyte differentiation and barrier function, placing it in a key position to control polarized epidermal functions downstream of the desmosome/IF network.

## Acknowledgments

We thank Alpha Yap, Andrew Kowalczyk, Lisa Godsel, and Quinn Roth-Carter for helpful feedback on the manuscript. Research reported in this publication was supported by Northwestern University Skin Biology & Diseases Resource-Based Center of the National Institutes of Health under award number P30AR075049. Imaging work was performed at the Northwestern University Center for Advanced Microscopy generously supported by NCI CCSG P30 CA060553 awarded to the Robert H Lurie Comprehensive Cancer Center. Multiphoton microscopy was performed on a Nikon A1R multiphoton microscope, acquired through the support of NIH 1S10OD010398-01. This work was supported by NIH grants R01 AR041836, R37 AR043380, with partial support from R01 CA228196, the J.L. Mayberry Endowment to K.J.G. and the Chicago Biomedical Consortium with support from the Searle Funds at The Chicago Community Trust. M.H. was supported by NIH F31 AR076188. J.A.B. was supported by NIH K01 AR075087 and T32 AR060710.

## Materials and methods

### Antibodies and reagents

The following primary antibodies were used: chicken anti-plakoglobin 1407 and 1408 (Aves Laboratories); rabbit anti-loricrin, rabbit anti-keratin 1, rabbit anti-keratin 5, and rabbit anti-keratin 14 (gifts from J. Segre, National Human Genome Research Institute); rabbit NW6 and NW161 anti-desmoplakin (Green laboratory); mouse DM1α anti-α-tubulin (Sigma-Aldrich, T6199); rabbit anti-Src (Sigma-Aldrich, HPA030875); rabbit anti-Fyn (Sigma-Aldrich, HPA023887); mouse anti-Yes (Santa Cruz Biotechnology, sc-46674); rabbit anti-GAPDH (Sigma-Aldrich, G9545); mouse JL8 anti-GFP (Clontech, 632381); mouse P124 anti-desmoglein 1 (Progen, 651111); mouse U100 anti-desmocollin 1a/b (Progen, 65192); mouse anti-vinculin (Sigma-Aldrich, V9131); mouse C4 anti-actin (EMD-Millipore, MAB1501); mouse HECD1 anti-E-cadherin (gift from M. Takeichi and O. Abe, Riken Center for Developmental Biology, Kobe, Japan); rabbit anti-EGF Receptor (4261), rabbit anti-phospho-EGF Receptor Tyr1068 (2234), rabbit anti-HER2/ErbB2 (2165), and anti-phospho-HER2/ErbB2 Tyr877 (2241) were from Cell Signaling Technology; mouse anti-ZO1 (BD BioSciences, 610967); rabbit anti-claudin 1 (51-9000), rabbit anti-claudin 4 (36-4800), and mouse anti-occludin (33-1500) were from Thermo Fisher Scientific. Secondary antibodies included goat anti-mouse, -rabbit, and -chicken HRP (Kirkegaard Perry Labs); goat anti-mouse, -rabbit and -chicken conjugated with Alexa Fluor 488, 568, or 647 nm (Thermo Fisher Scientific, 1:300 IF); goat anti-mouse, -rabbit, and -chicken conjugated with Alexa Fluor 488, 568, or 647 nm (Thermo Fisher Scientific); goat anti-mouse IgG1 isotype conjugated with Alexa Fluor 488, 568, or 647 nm (Thermo Fisher Scientific); and goat anti-mouse IgG2a isotype conjugated with Alexa Fluor 488, 568, or 647 (Thermo Fisher Scientific). Alexa Fluor 488, 568, or 647 nm phalloidin (Thermo Fisher Scientific) was used to stain filamentous actin and 4′,6-Diamido-2-Phenylindole (DAPI, Sigma-Aldrich) to stain nuclei.

### Western blot analysis

Whole cell lysates were generated using Urea Sample Buffer (8 M urea, 1% SDS, 60 mM Tris, pH 6.8, 5% β-mercaptoethanol, 10% glycerol). Proteins were separated by SDS-PAGE electrophoresis and transferred to nitrocellulose membranes. Membranes were blocked with either 5% milk or 1% BSA in PBS or TBS with or without 0.05% Tween. Primary and secondary antibodies were incubated in this blocking solution. Immunoreactive proteins were visualized using chemiluminescence. Densitometry analysis was performed using ImageJ.

### Cell culture

Primary normal human epidermal keratinocytes (NHEKs) were isolated from neonatal foreskin provided by the Northwestern University Skin Biology and Disease Resource-based Center (SBDRC) as previously described [67]. Cells were maintained in growth medium (M154 media supplemented with 0.07 mM CaCl_2_, human keratinocyte growth supplement (HKGS), and gentamicin/amphotericin B solution (G/A); Thermo Fisher Scientific). NHEKs were induced to differentiate by addition of 1.2 mM CaCl_2_ to growth medium with or without addition of 1 μg/mL ephrin A1-FC peptide (R&D Systems) for the indicated times. NHEKs were used to generate 3D epidermal equivalent cultures as previously described previously [67]. Pharmacological treatments included 1 unit/ml Rho Activator I (CN01; Cytoskeleton, Denver, CO), 1 mM TAK165 (Mubritinib, Selleck Chemicals), dimethyl sulfoxide (DMSO, Sigma-Aldrich), 20 μM PP2 (Sigma-Aldrich), and 60 nM CCG-1423 (Sigma-Aldrich).

NHEKs were transfected with either siGENOME Non-Targeting siRNA Pool #2 (siCtl) or siGENOME SMARTpool siRNA D-019800-17 DSP (Dharmacon, Lafayette, CO) (siDP1) or stealth siRNA oligonucleotides 5′-CAGGGCUCUGUCUUCUGCCUCUGAA-3′ from Life Technologies (siDP2) using the Amaxa Nucleofector System (Lonza) according to manufacturer’s instructions. NHEKs were suspended in Ingenio Electroporation Solution (Mirus) with siRNA (final concentration 50 nM) and electroporated using program X-001.

### Viral transduction

DPNTP-GFP was cloned from pEGFP-N1-DPNTP into pLZRS using the standard Gateway/TOPO cloning protocol. pLZRS-miR Dsg1 (shDsg1) was generated as previously described (Getsios et al., 2009). LZRS–NTshRNA (shCtl) was generated with the following sequences inserted: NTshRNA-fwd 5′–GTATCTCTTCATAGCCTTAAA–3′ and NTshRNA-rev 5′–TTTAAGGCTATGAAGAGATAC–3′. Keratinocytes were transduced with retroviral supernatants produced from Phoenix cells (provided by G. Nolan, Stanford University, Stanford, CA) as previously described (Simpson, Kojima, & Getsios, 2010). Briefly, Phoenix cells transiently transfected with retroviral pLZRS cDNA constructs were harvested at 70% confluency for 24 hours at 32°C. Supernatants were collected and concentrated using Centricon Plus-20 columns (EMD Millipore). Infection of keratinocytes was done at 15% cell confluence with incubation at 32°C for 1.5 hours in M154 media containing 4 μg/ml polybrene (Sigma-Aldrich, H9268) and retrovirus supernatants. Lentivirus (pLVX myristoylated tdTomato, Clontech) transduction of keratinocytes utilized virus obtained from the Northwestern University SBDRC. Briefly, keratinocytes were plated at 30% confluence and the next day treated with 1 μg/ml polybrene plus lentivirus and incubated at 37°C overnight followed by washing

### Immunofluorescence/Microscopy

For immunofluorescence analysis, NHEKs cultured on glass coverslips were fixed either in anhydrous ice-cold methanol for 3 min on ice or 4% paraformaldehyde solution for 15 min at room temperature. Cells were permeabilized and blocked with 0.25% Triton X-100 and 5% goat serum for 30 minutes and then processed for immunofluorescence. For skin biopsies, paraffin-embedded sections were baked overnight at 60°C and deparaffinized using xylene/ethanol. After permeabilization with 0.5% Triton X-100 in PBS, antigen retrieval was performed using 0.01 M citrate buffer, pH 6.0. All samples were mounted onto glass slides or using glass coverslips with ProLong Gold antifade reagent (Thermo Fisher).

Apotome images were acquired using ZEN 2.3 software (Carl Zeiss) with an epifluorescence microscope system (Axio Imager Z2, Carl Zeiss) fitted with an X-Cite 120 LED Boost System, an Apotome.2 slide module, Axiocam 503 Mono digital camera, and a Plan-Apochromat 40x/1.4 or Plan-Apochromat 63x/1.4 objective(Carl Zeiss). Confocal z-stacks (z-step size of 0.24-0.5 μm) were acquired using a Nikon A1R confocal laser microscope equipped with GaAsP detectors or a Nikon W1 Spinning Disk Confocal with a 95B prime Photometrics camera and a 60× Plan-Apochromat objective lambda with a NA of 1.4 and run by NIS-Elements software (Nikon). NIS-Elements (version 5.02) was used to generate 3D reconstructions of z-stacks using the Volume Viewer tool with z-depth coding blending (rainbow contrast lookup table) or ImageJ was used to generate color coded z-projections using Temporal-Color Code (Fire lookup table). All fluorescence intensity-based quantification was performed using ImageJ.

### Laser ablation

NHEKs were transduced with pLVX myristoylated tdTomato (myr-tomato) lentivirus to allow tracking cell outlines and 24 hours after they were transduced with either pLZRS GFP or DPNTP-GFP retrovirus. These cells were then used to generate 3D epidermal equivalent cultures as described above. At the indicated timepoints, sufficient medium was added to a culture dish to re-submerge the epidermal equivalent to perform imaging with a water dipping objective and secured using a harp anchor. The culture was allowed to equilibrate for 20 minutes before imaging and only imaged for 1 hour each. Two-photon laser ablation was used to assess intercellular forces. Briefly, ablation was performed on a Nikon A1R-MP+ multiphoton microscope running Elements version 4.50 and equipped with an Apo LWD 25× 1.10W objective. Cells were maintained at 37 °C and 5% CO_2_. Images of myr-tomato and GFP were obtained at a rate of 1 frame per second for 2 s before and 45 s after ablation using 4% laser power at 970 nm. Ablation was performed using 40% laser power at a scan speed of 512. The distance between the cell–cell vertices over time was measured using the Manual Tracking plugin in ImageJ. Distance curves were then generated using Excel software. Initial recoil measurements were calculated using Prism 8 software as previously described [68].

### Sorting/stratification assay

Wild type NHEKs were mixed at known ratios with genetically-modified NHEKs that were transduced with GFP, DPNTP-GFP, Dsg1-GFP, and in combination as indicated with subsequent nucleofection with siRNA. These combined populations were switched to M154 medium containing 1.2 mM calcium, HKGS, and G/A. Cultures were allowed to stratify for 3 days before fixation using 4% paraformaldehyde. For Dsg1-GFP experiments, cultures were allowed to stratify for 1 day before fixation with 4% paraformaldehyde. Samples were processed for immunostaining as above for plakoglobin to visualize all cell outlines as well as DAPI to visualize nuclei. Confocal z-stacks were obtained and processed in ImageJ to contain either only basal cells or suprabasal cells by selecting a subset of z-frames. The total number of cells per field was the quantified using DAPI and plakoglobin staining. The subset of the total population that expressed GFP was then calculated and compared to the percentage of cells predicted to express GFP based on the number of wild type and GFP expressing cells put into the system. This was compared for basal as well as suprabasal cells.

### Transepidermal electrical resistance

Costar 24 mm Transwell 0.4 μm pore inserts (Costar, 3450) were coated with CELLstart CTS Substrate (Thermo Fisher Scientific, A10142-01) according to manufacturer. NHEKs were seeded at confluence in Defined PCT Epidermal Keratinocyte Medium, (CnT-07, Zenbio) and submerged in CnT-07 for 2 days with medium in both upper and lower chambers. On the third day, CnT-07 was replaced with CnT-Prime 3D Barrier Culture Medium (CnT-PR-3D, Zenbio) with cells submerged. The following day, “Day 0” TEER measurements were taken and medium was removed from the upper chamber to expose NHEKs to an air-liquid interface. Subsequent TEER measurements were taken every other day/ at indicated times by adding pre-warmed (37°C) CnT-3D-PR to the top chamber. Measurements were made with an EVOM epithelial volt/ohm meter (World Precision Instruments, Sarasota, FL, USA). Measurements were taken in triplicate per sample and averaged. Medium was removed promptly from the top chamber as soon as TEER measurements were taken.

### Statistical Analysis

For all assays, at least three independent experiments were carried out and exact numbers are found in the figure legends. Independent experiments are shown in corresponding colors in figure graphs. Independent experiments were considered as those performed with NHEKs derived from individual patient materials. Data in all graphs are presented as means and error bars represent standard error of the mean (SEM). Graphs were generated using Excel or Prism software. Statistical analyses were performed using Prism software and are specifically indicated in figure legends. P values less than 0.05 were considered statistically significant.

---

**Figure S1.**
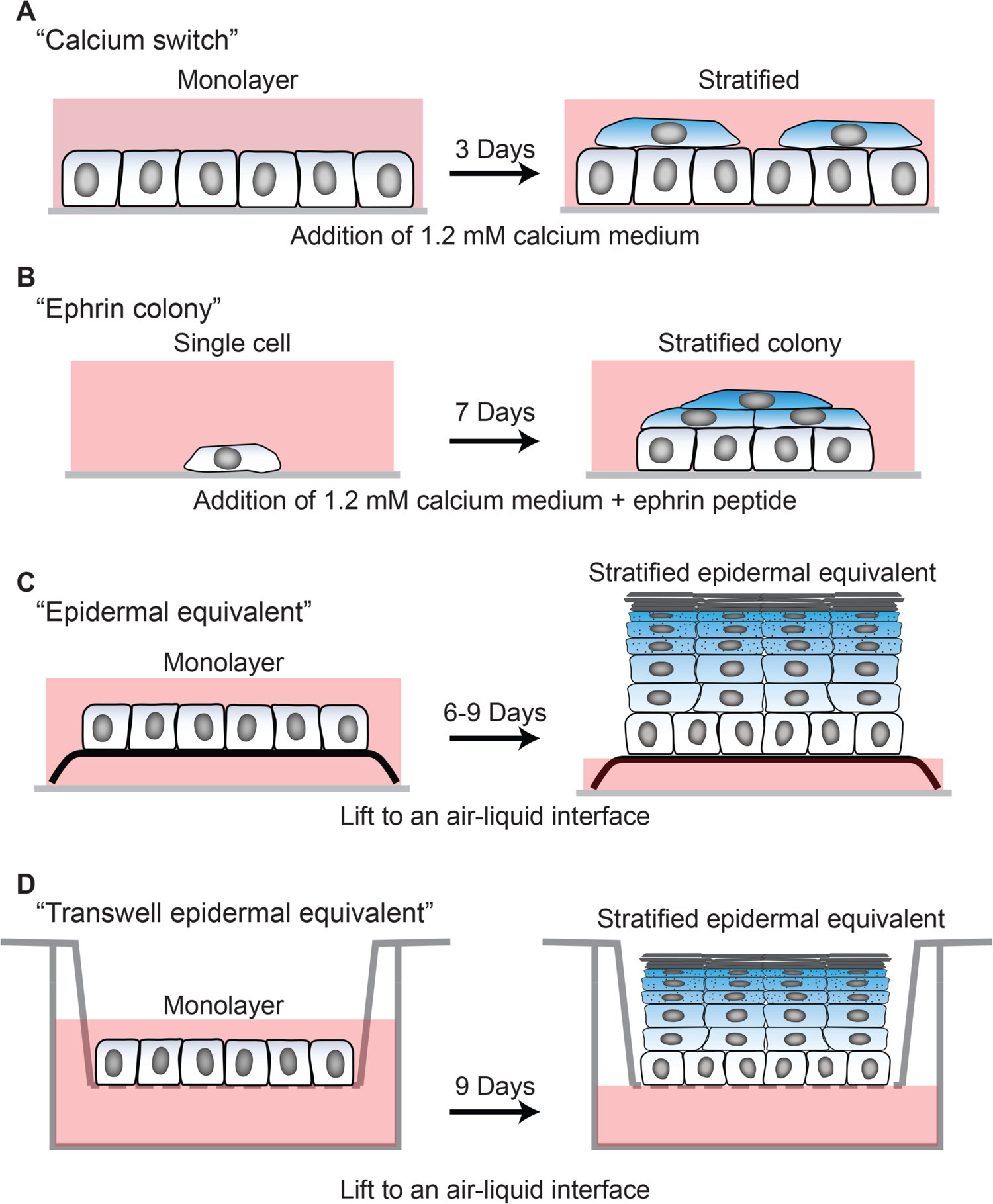
Cell culture models used in this study. A) Calcium switch experiment-NHEKs are seeded at confluence in 0.07 mM calcium containing medium. The following day, the medium is changed to 1.2 mM calcium containing medium to induce stratification and differentiation. After 3 days, unless otherwise noted, cells were harvested for analysis. B) Ephrin colony experiment-NHEKs are seeded at low confluence such that they are initially sparse fields of single cells that subsequently grow into individual colonies. Culture medium contains 1.2 mM calcium and 1 μg/ml ephrin-A1 peptide to induce stratification and differentiation. After 7 days, unless otherwise noted, cells were harvested for analysis. C) Epidermal equivalent experiment-monolayers of NHEKs were seeded on fibroblast containing collagen I gels at confluence. Subsequently, the gels were lifted to an air-liquid interface by placing them onto a metal grid with culture medium only filling the space up to the bottom of the metal grid. Cultures were allowed to stratify and differentiate for 6-9 days prior to harvesting, unless otherwise noted. D) Transwell epidermal equivalent experiment-monolayers of NHEKs were seeded onto transwell inserts at confluence. The cultures were exposed to an air liquid interface by removing the medium from the top chamber of the transwell. Cultures were allowed to stratify and differentiate for 9 days prior to harvesting, unless otherwise noted. Transepidermal electrical resistance measurements were performed by adding medium into the top chamber, taking measurements, then promptly removing medium to restore the air-liquid interface.

**Figure S2.**
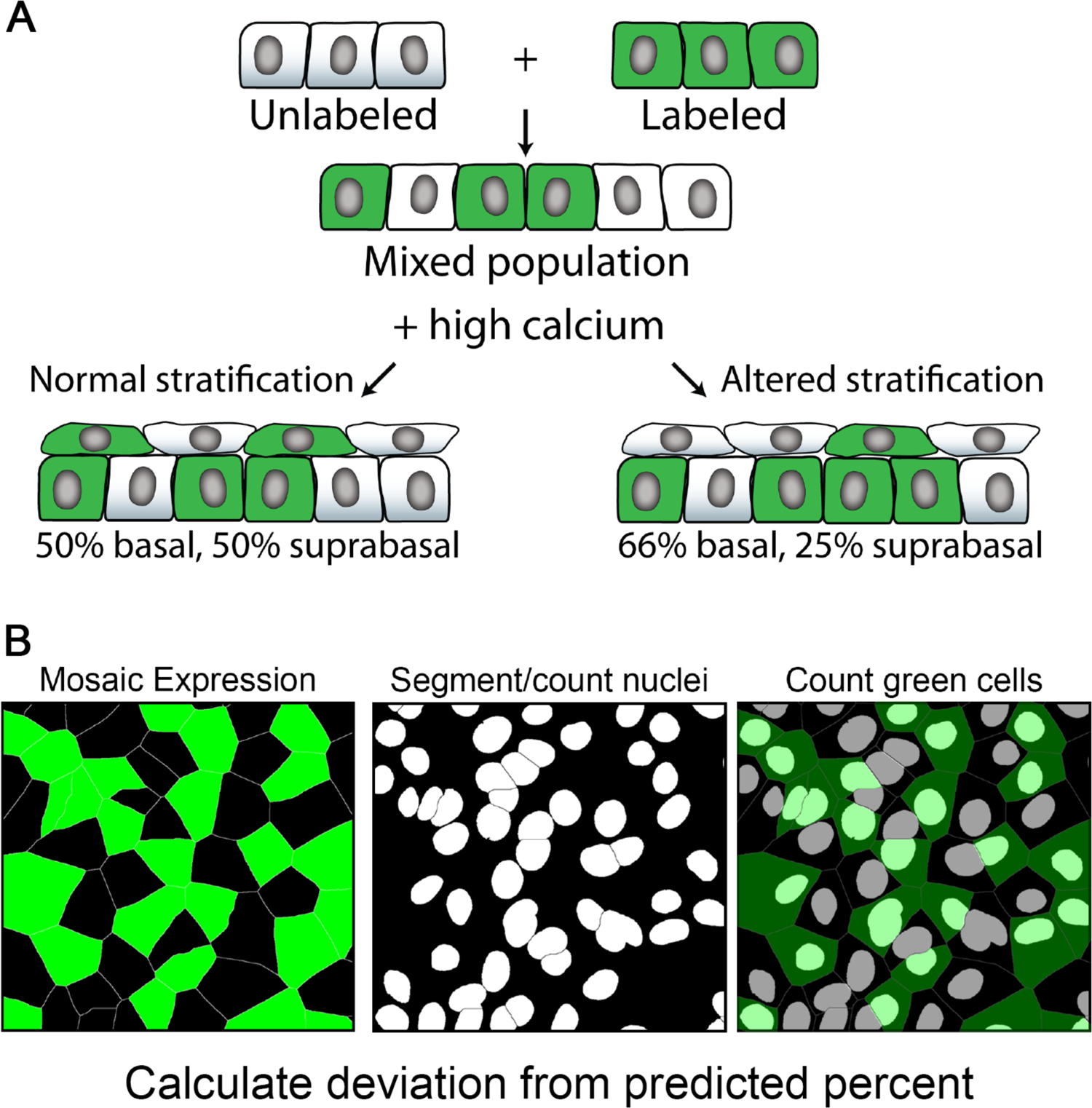
Schematic of stratification assay. A) Wild type NHEKs were mixed at known ratios with genetically-modified NHEKs that expressed GFP. These combined populations were switched to 1.2 mM calcium (high calcium) containing medium and allowed to stratify for 3 days. An example in which 50% of cells are expressing GFP and 50% are wild type is presented. If stratification occurs normally, 50% of both basal and suprabasal cells will be expressing GFP. If stratification is inhibited, there will be an underrepresentation of GFP-positive cells in the suprabasal layer. B) To quantify the percentage of cells expressing GFP, total cells were segmented (using a nuclear dye for example) and the percentage of GFP positive cells was calculated and then compared to the predicted percentage based on the initial ratio of cells plated.

**Figure S3.**
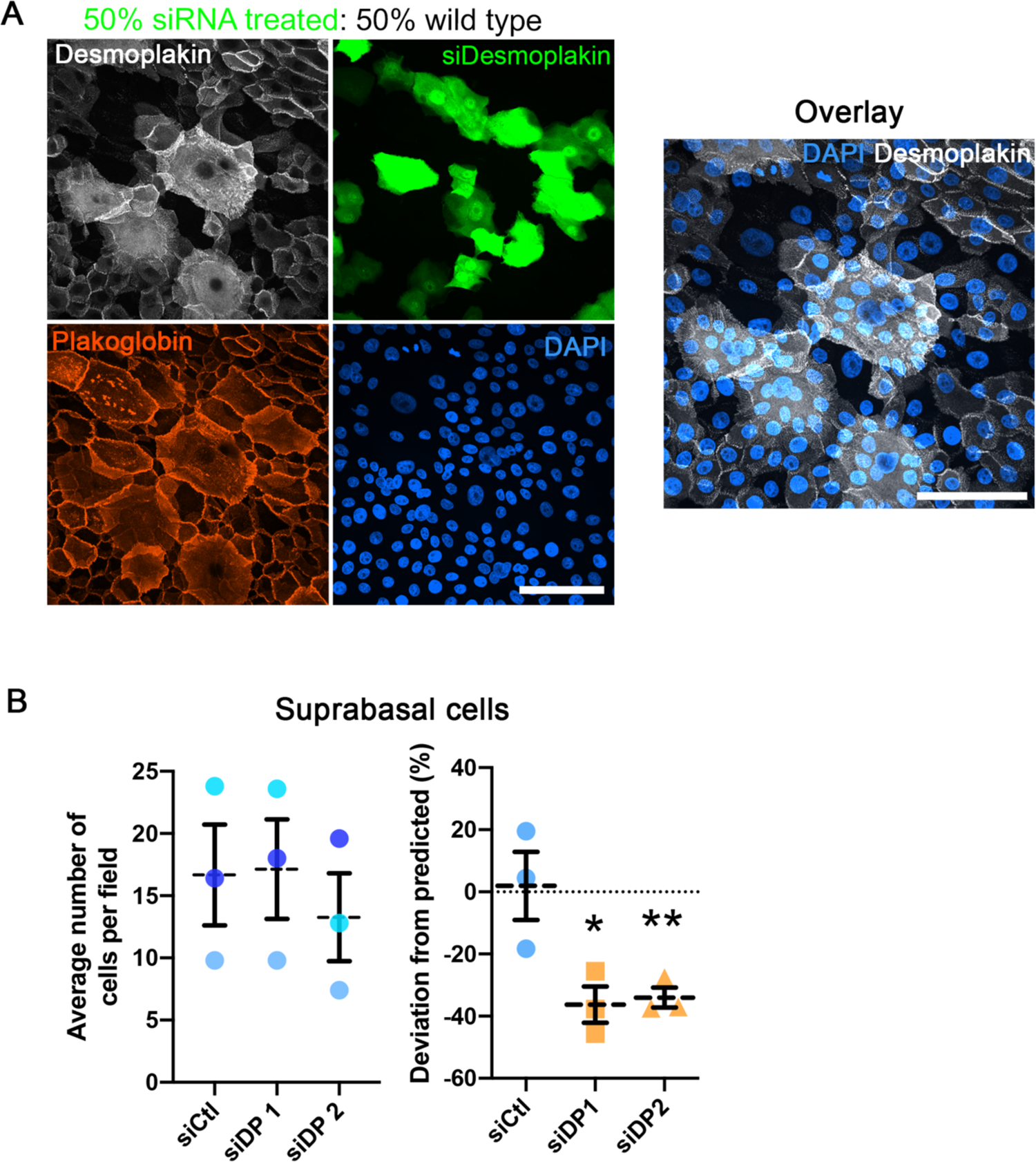
RNAi-mediated depletion of desmoplakin impairs stratification. A) A representative sorting assay for desmoplakin (DP) depleted cells is shown. GFP-positive cells treated with either non-targeting (siCtl) or two DP-targeting siRNA pools (siDP) were mixed with wild type cells and induced to stratify. Immunostaining for DP indicates level of knockdown and the percentage of suprabasal cells (total cells identified with plakoglobin in red and DAPI in blue) that were GFP positive was calculated. Bar is 100 μm. B) Quantification for the average number of suprabasal cells as well as the deviation from the predicted percentage of GFP positive cells in the suprabasal layer for control (siCtl) and DP-depleted (siDP1 and siDP2) conditions is shown. Dashed lines indicate the mean of 3 independent experiments and error bars are SEM. *p=0.025, **p=0.0087, one sample t test with theoretical mean of 0.

**Figure S4.**
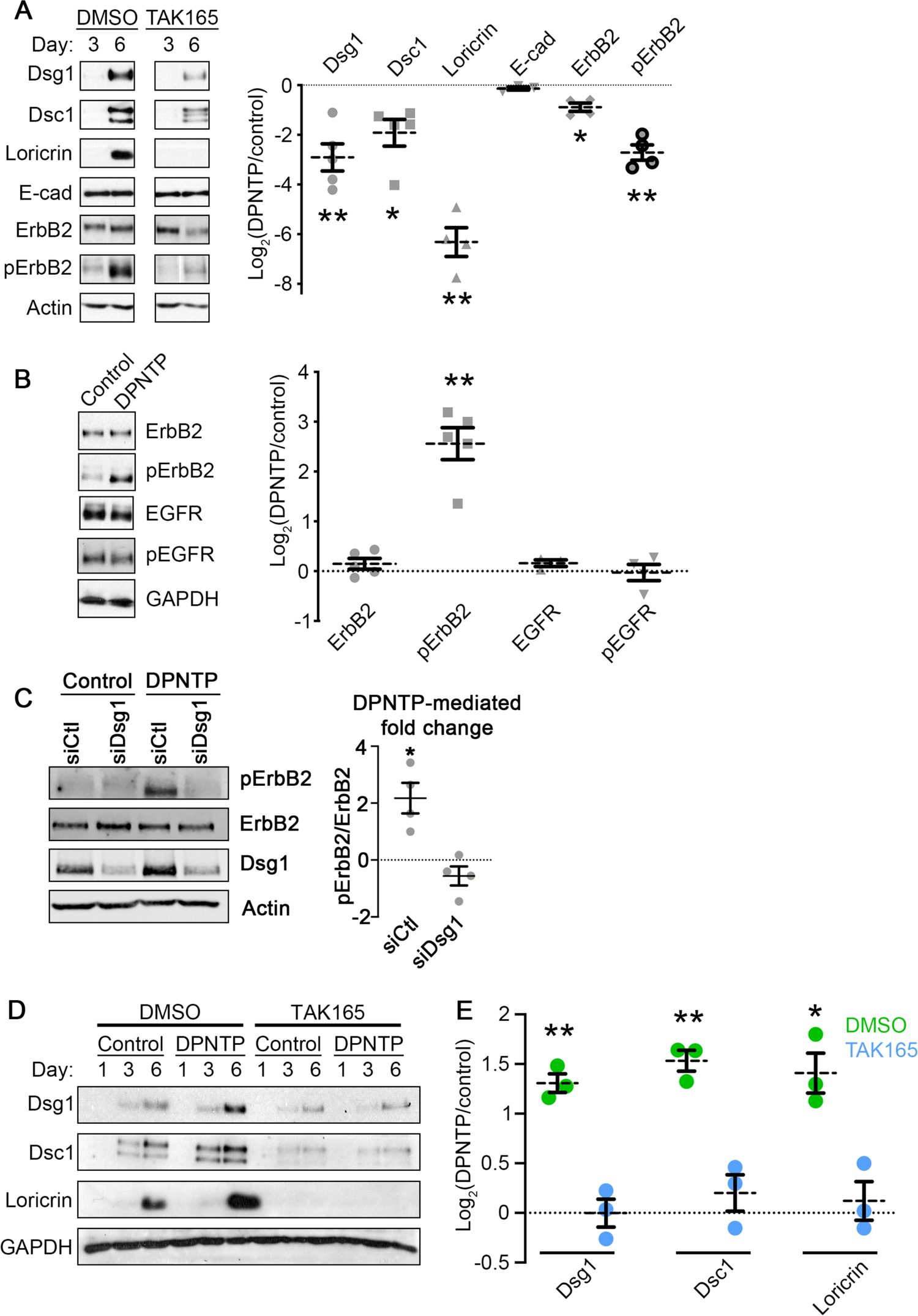
The DPNTP-mediated effects on differentiation are dependent on ErbB2. A) Left, western blots show the expression of the indicated proteins in control (DMSO) or ErbB2 inhibitor (TAK165) treated keratinocytes at the indicated times after addition of 1.2 mM calcium medium. Right, quantification of the fold change (Log_2_-transformed) of TAK165 treated over control are shown for the indicated protein expression. Dashed lines indicate the mean of 3-5 independent experiments and error bars are SEM. *p*≤*0.02, **p*≤*0.006, one sample t test with theoretical mean of 0. B) Left, western blots show the expression of the indicated proteins in GFP (Control) or DPNTP-GFP expressing keratinocytes differentiated in 1.2 mM calcium medium for 6 days. Right, quantification of the fold change (Log_2_-transformed) of DPNTP-expressing over control are shown for the indicated protein expression. pErbB2 is Y877-phosphorlated ErbB2 and pEGFR is Y1068-phosphoylated EGFR. Dashed lines indicate the mean of 3-5 independent experiments and error bars are SEM. **p=0.0013, one sample t test with theoretical mean of 0. C) Left, western blots show the expression of the indicated proteins in GFP (Control) or DPNTP-GFP expressing keratinocytes treated with the indicated siRNA and differentiated in 1.2 mM calcium medium for 6 days. Right, quantification of the fold change (Log_2_-transformed) of DPNTP-expressing over control are shown for the indicated protein expression. pErbB2 is Y877-phosphorlated ErbB2. Means are from 3 independent experiments and error bars are SEM. **p=0.0385, one sample t test with theoretical mean of 0. D) Western blots show a time course of the expression of the indicated proteins in differentiated GFP (Control) or DPNTP-GFP expressing keratinocytes either treated with DMSO as a control or the ErbB2 inhibitor TAK165. E) Quantification of the fold change (Log_2_-transformed) of DPNTP-expressing over GFP control keratinocytes are shown for the indicated protein expression for both DMSO and TAK165 treatment. Dashed lines indicate the mean of 3 independent experiments and error bars are SEM. *p=0.02, **p*≤*0.05, one sample t test with theoretical mean of 0.

**Figure S5.**
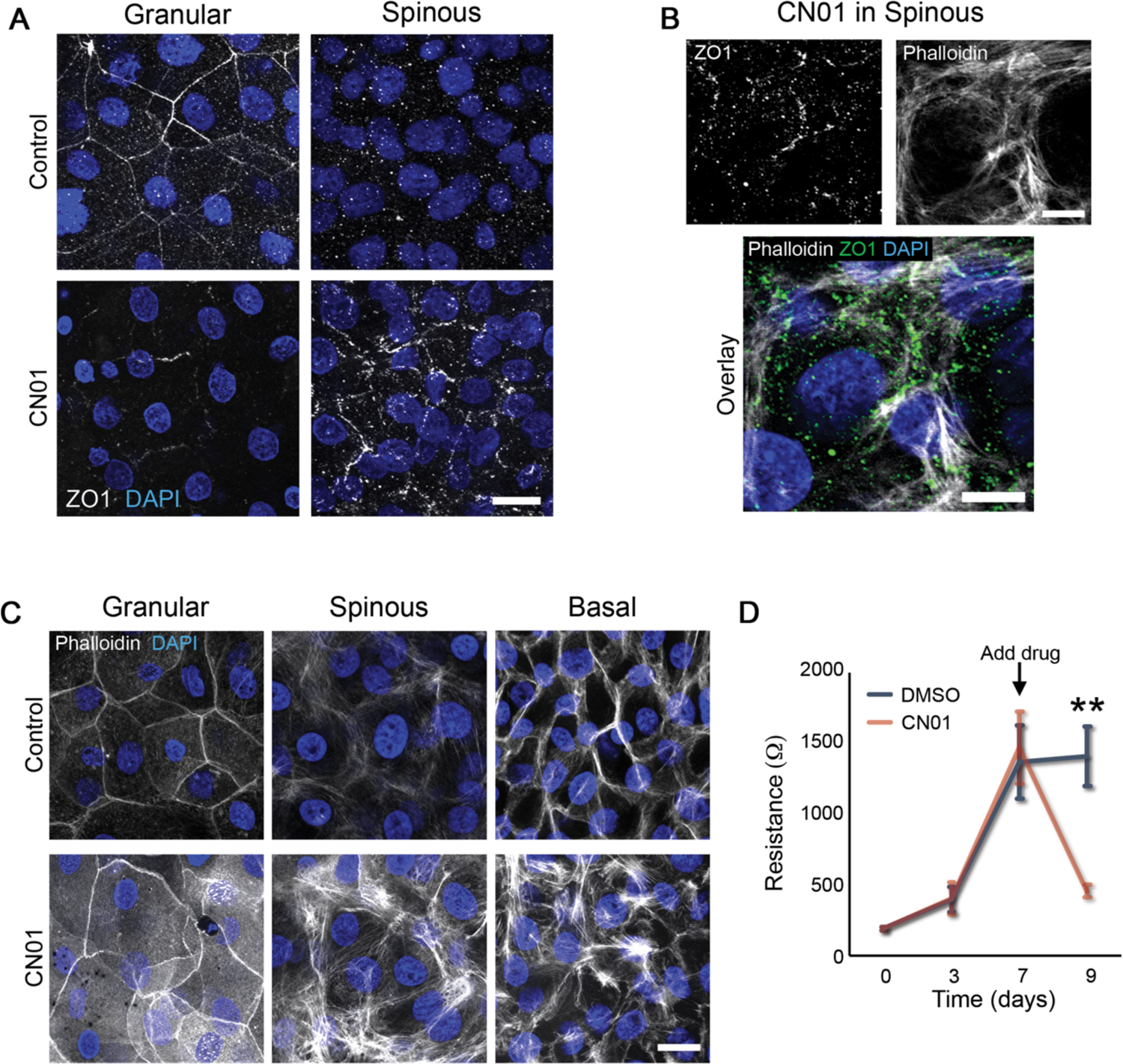
Excessive actomyosin contraction impairs epidermal barrier function. A) Maximum projection micrographs of day 9 transwell epidermal equivalent cultures that were treated with either DMSO (Control) or a Rho activator (CN01, 1 unit/mL) at day 7-9 show immunostaining for the tight junction protein ZO1 and staining for nuclei in blue (DAPI) within the indicated cell layers. Bar is 20 μm. B) Maximum projection micrographs of day 9 transwell epidermal equivalent cultures that were treated with a Rho activator (CN01, 1 unit/mL) at day 7-9 show immunostaining for the tight junction protein ZO1, staining for nuclei in blue (DAPI), and F-actin (phalloidin) within the spinous cell layer. Bar is 10 μm. C) Maximum projection micrographs of day 9 transwell epidermal equivalent cultures that were treated with either DMSO (Control) or a Rho activator (CN01, 1 unit/mL) at day 7-9 show staining for nuclei in blue (DAPI) and F-actin (phalloidin) within the indicated cell layers. Bar is 20 μm. D) Quantification of resistance measurements from TEER experiments performed on a time course of transwell epidermal equivalent cultures that were treated with either DMSO or a Rho activator (CN01, 1 unit/mL) at day 7. Data are presented as mean ± SEM. **p=0.0029, paired t test from 6 independent experiments.

**Figure S6.**
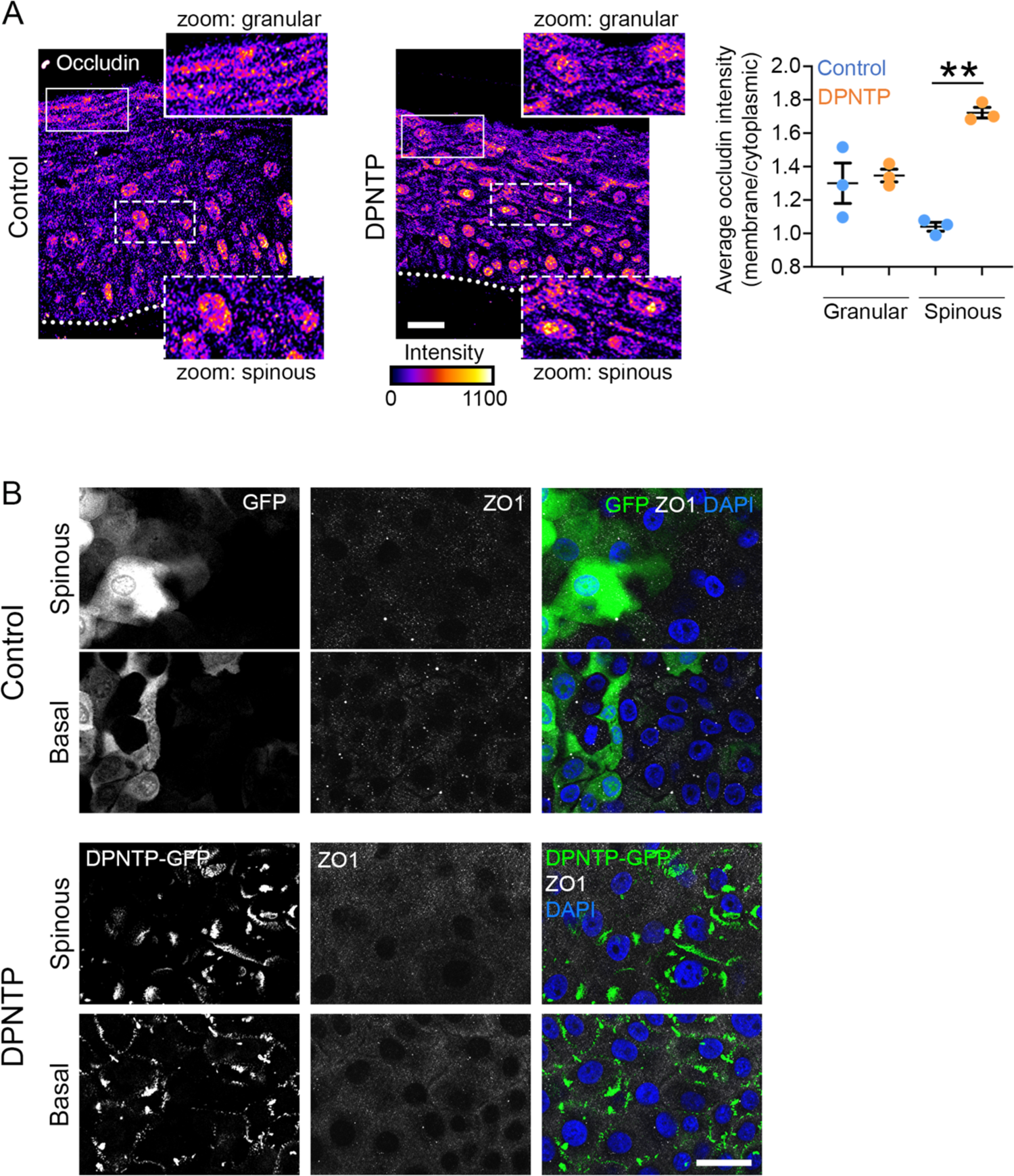
Expression of DPNTP and layer-specific tight junction component staining. A) Left, micrographs of transverse sections of day 6 human epidermal equivalent cultures expressing either GFP (Control) or DPNTP-GFP immunostained for the tight junction component occludin are shown. Dotted line marks the bottom of the basal layer. Bar is 20 μm. Intensity is displayed with the indicated lookup table. Right, quantification of the average membrane to cytoplasmic ratio of occludin staining in the indicated cell layers is shown. Means are from 3 independent experiments and error bars are SEM. **p=0.0014, paired t test. B) Fluorescence micrographs show the expression of GFP (Control) and DPNTP-GFP in the spinous and basal layer of day 9 transwell epidermal equivalent cultures (same as Figure 4D) that were immunostained for the tight junction component ZO1. Nuclei are stained with DAPI in blue. Bar is 25 μm.

**Figure S7.**
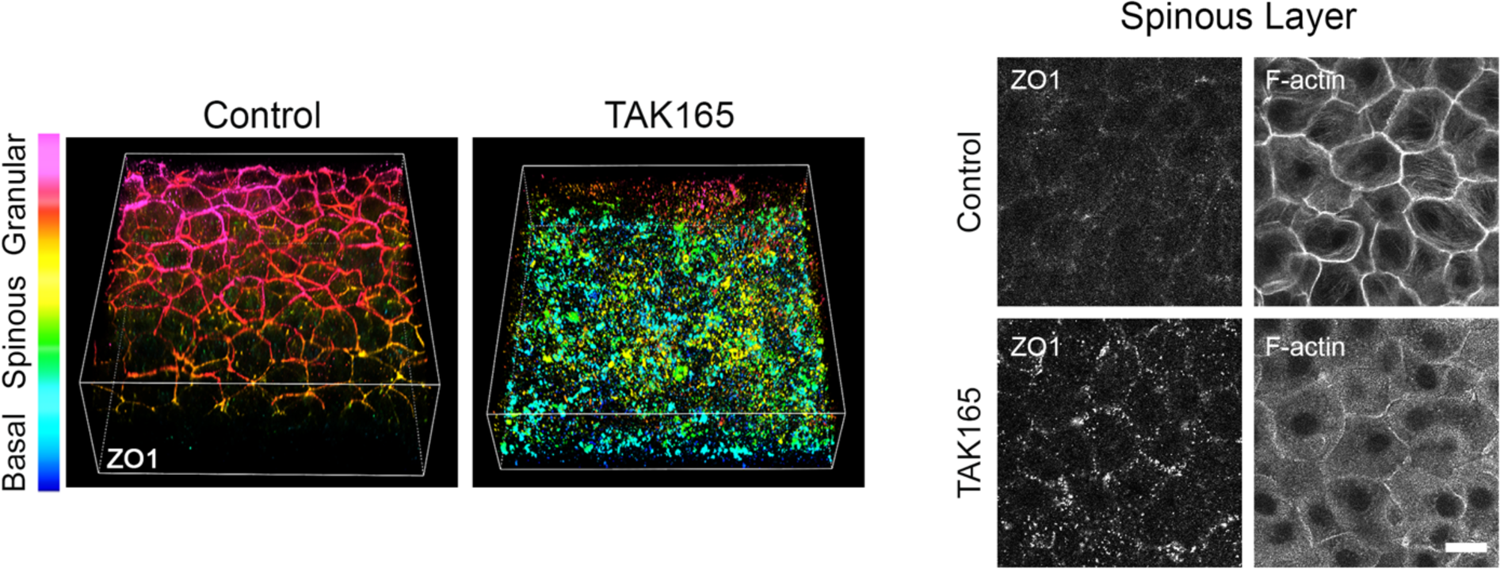
Treating epidermal equivalent cultures with an ErbB2 inhibitor disrupts polarized ZO1 immunostaining. 3D renderings of whole mount immunostaining of tight junction component ZO1 in control (DMSO) and TAK165 treated day 9 epidermal equivalent cultures are presented with a look-up table that depicts z-depth in the indicated colors. Right panels show representative ZO1 immunostaining and F-actin (phalloidin staining) in the spinous layer. Cultures were allowed to grow normally until day 7 and then treated with DMSO or TAK165 for 2 subsequent days. Bar is 20 μm.

Movie S1. Tension in suprabasal layer. A representative timelapse of a laser ablation experiment in the suprabasal layer in a Day 6 3D epidermal equivalent culture. Cell outlines are visualized with myr-tomato and time between frames is 1 sec. Total time is 45 sec. Field of view is 79.5×79.5 μm.

